# Phylogenomics and Fossilized Birth-Death Dating Reveals Extensive Post-Cretaceous Worldwide Diversification of Cicadidae (Hemiptera, Auchenorrhyncha)

**DOI:** 10.1101/2025.11.08.687401

**Authors:** Mark Stukel, Jordan Douglas, Tatiana Petersen Ruschel, Stéphane Puissant, Ben W. Price, Martin Villet, Alan R. Lemmon, Emily Moriarty Lemmon, Chris Simon

## Abstract

The Cretaceous-Paleogene (K-Pg) mass extinction resulted in a massive turnover in biodiversity. The oldest fossils of the globally-distributed insect family Cicadidae date to the Paleocene, suggesting that this family diversified after the K-Pg boundary. We analyzed 490 nuclear Anchored Hybrid Enrichment loci as well as mitochondrial genomes for the Cicadidae, sampling all five subfamilies and 82% of tribes, using concatenated maximum-likelihood and multi-species coalescent approaches to resolve the phylogenetic relationships of Cicadidae subfamilies. We estimated divergence times of Cicadidae lineages using a fossilized birth-death model augmented into the multispecies coalescent with 44 fossil taxa. We estimated a Cretaceous origin for Cicadidae with four of the five subfamilies diversifying shortly after the K-Pg extinction event. Our fossilized birth-death tip-dating approach improved the precision of age estimates for many cicada clades compared to dates based on node-calibration from previous studies. Our results augment insight into how the K-Pg mass extinction shaped present-day diversity.

## Introduction

The timing of diversification events are important for understanding processes that generate biodiversity. Throughout geological time, mass extinctions played a crucial role in shaping biodiversity through accelerated extinction and subsequent diversification. The most recent mass extinction event, the Cretaceous-Paleogene (K-Pg) transition approximately 66 million years ago involving the extinction of around 76% of all species, helped to shape which evolutionary lineages are most diverse today (Schulte et al. 2010). The mechanism behind post-extinction diversification is thought to be competitive release, when lineages surviving mass extinctions face much less ecological competition and rapidly radiate (Erwin 2001; Jablonski 2001; Hull 2015; Hoyal Cuthill et al. 2020). Understanding mass extinction events and their associated evolutionary radiations provides insight into the historical development of biodiversity and the processes that may shape future biodiversity.

Phylogenetic tree inference for evolutionary radiations is challenging. Studies using a handful of genes are frequently sufficient to infer phylogenetic trees for taxa that underwent well-spaced speciation events; however, these genes often do not have enough phylogenetic information to resolve the short branches that are typical of episodic evolutionary radiations (Whitfield and Kjer 2008). Furthermore, the assumption implicit in analyses of concatenated sequences is violated for most datasets – i.e., that individual genes all share the same evolutionary history as the species tree (Degnan and Rosenberg 2006; Salichos and Rokas 2013). Concatenated sequences can therefore lead to biased estimates of substitution rates and divergence times (Kubatko et al. 2011; Mendes and Hahn 2016; Ogilvie et al. 2016), and inconsistently estimate the tree topology (Heled and Drummond 2010; Ogilvie et al. 2017). Avoiding these biases necessitates the use of the multispecies coalescent (MSC) to create species trees from non-concatenated genes (Maddison 1997; Degnan and Rosenberg 2009; Edwards 2009; Liu et al. 2009). Genomic datasets and analytical methods that account for deep coalescence (also called incomplete lineage sorting, or ILS) have shown their utility for phylogenetic problems such as evolutionary radiations (Hime et al. 2020; Owen and Miller 2022; Smith et al. 2023; Stukel et al. 2024).

Challenges also exist in dating phylogenomic data. Bayesian fossilized birth-death dating has the advantage over node dating in that prior calibration densities are replaced with a model incorporating paleontological information into the diversification process (Heath et al. 2014). It also allows all available fossils to be incorporated into the estimate rather than simply the oldest fossil assigned to a node. Unfortunately, Bayesian molecular dating methods using the MSC are too computationally intensive to handle hundreds of genes and many taxa simultaneously. Some strategies for reducing the computational complexity of dating large data sets include: 1) fixing the maximum-likelihood tree topology and estimating divergence times using a subset of clock-like genes (Smith et al. 2018); 2) estimating divergence times using relaxed clock models in a maximum-likelihood framework (Smith and O’Meara 2012; Tamura et al. 2012); 3) dividing a phylogeny into a backbone tree and substantiated subtrees for separate inference, followed by rejoining the subtrees to the backbone tree; 4) or using multiple smaller samples of genes and assessing whether they produce similar species phylogenies. Here we employ the third and fourth strategies to date the origin of cicadas in relation to the K-Pg boundary in a Bayesian framework.

### Study system

Cicadas are an excellent model lineage of insects for understanding evolutionary radiations. Cicadoidea contains two families: Tettigarctidae (hairy cicadas) and Cicadidae (singing cicadas) (Moulds 2005). Cicadidae is distributed globally (except for Antarctica) with over 3,400 described species, while Tettigarctidae, which was palaeontologically a Eurasian lineage (Jiang et al. 2024), is now restricted to the southeastern Australian mainland and Tasmania with two extant species (World Auchenorrhyncha Database, accessed 18 April 2025). Cicadidae comprises five subfamilies: Cicadinae and Cicadettinae, both with global distributions; Tibicininae, distributed across the Neotropics, Holarctic, and North Africa (Puissant and Gurcel 2023); Tettigomyiinae, in Africa and Madagascar; and the monogeneric Neotropical Derotettiginae (Marshall et al. 2018; Simon et al. 2019; Owen et al. 2022). These subfamilies house 55 tribes and over 500 genera (World Auchenorrhyncha Database, accessed 18 April 2025). Due to their species-specific songs, cicadas have been considered a model system for studying speciation and reproductive isolation (Villet 1995; Marshall and Cooley 2000; Marshall et al. 2011; Hertach et al. 2016). Furthermore, due to their long underground life cycles, short-lived adults, and low dispersal ability (Tobella et al. 2025), they have clear population genetic structure that has been analyzed to answer biogeographic and phylogeographic questions (e.g., Hill et al. 2009, 2021; Marshall et al. 2009, 2012; Seabra et al. 2009; Price et al. 2010, 2019; Hertach et al. 2016; Bator et al. 2021; Nguyen et al. 2022; Stukel et al. 2024).

The fossil record suggests that Cicadidae may have originated or diversified around the time of the K-Pg mass extinction, between the Late Mesozoic and Early Cenozoic. Although all fossils unambiguously assigned to crown Cicadidae are from the Cenozoic (Moulds 2018; Moulds et al. 2022, 2023; Boderau et al. 2025b, 2025a), a recent morphological phylogenetic study has suggested that some Mesozoic fossils traditionally assigned to Tettigarctidae are in fact fossils branching off the stem lineage of Cicadidae (Jiang et al. 2024). At the same time fossils assigned to Tettigarctidae appear as early as the Triassic period (Jouault et al. 2024), are numerous throughout the Mesozoic with much of the diversity found in the Jurassic, but decline in the Cenozoic (Shcherbakov 2009; Moulds 2018). This timeline of fossils raises the possibility that the K-Pg extinction event was an important factor for diversification of the Cicadidae.

Previous studies have investigated multiple evolutionary radiations at different taxonomic levels within the Cicadidae. These include a recent (ca. 14 Ma) New Zealand cicada radiation of five genera and around 60 species (Arensburger et al. 2004; Buckley et al. 2006; Buckley and Simon 2007; Marshall et al. 2008, 2011, 2012; Bator et al. 2021). Other radiations at deeper levels include tribe-level radiations, such as of the Tacuini (Hill et al. 2015) with over 200 described species distributed in North America, Europe, Asia, SE Asia, Oceania, Australia, and northern South America, the Platypleurini with over 250 described species distributed from South Africa to Korea (Price et al. 2019), and the Cicadettini with almost 600 described species and a global distribution except Madagascar (Marshall et al. 2016). Other cicada radiations include collections of tribes, such as a radiation of over 10 Asian cicada tribes with over 700 described species (Hill et al. 2021), or the entire family (Owen et al. 2022). While many previous studies investigating these cicada radiations at different taxonomic levels using small numbers of genes were able to resolve some relationships among taxa, many areas of the phylogeny remained unresolved, requiring phylogenomic approaches to improve our understanding of the evolutionary history of cicadas. In particular, Owen et al. (2022) dramatically improved the phylogenetic resolution of the entire family, but their study included several undersampled lineages that are better represented in the present study.

We had two main aims for this study. First, we resolve the evolutionary relationships for Cicadidae using a phylogenomic dataset with expanded taxon sampling compared to Owen et al. (2022), with 60% more taxa comprising one additional subfamily, three additional tribes, and 37 additional genera and to increase the sampling for South America, Africa, and Madagascar, improving the geographic breadth and evenness for the taxon sampling. Second, we investigate whether the K-Pg extinction event was associated with Cicadidae diversification by inferring divergence times under an MSC framework using the same phylogenomic dataset.

## Methods

Data and scripts for all analyses described below are deposited on Dryad:

### Taxon sampling

We expanded the taxon sampling of a previous phylogenomic dataset for Cicadidae (Owen et al. 2022), with focus on subfamilies and tribes that were under sampled, especially Afrotropical and Neotropical taxa. This allowed our expanded dataset to better span the phylogenetic and biogeographical diversity of Cicadidae. Cicada specimens were primarily collected by members of the Simon lab or by specialist collaborators, with the exception of six specimens that were borrowed from the Natural History Museum, London and one specimen loaned by the Muséum national d’Histoire naturelle, Paris (Table S1). Cicada specimens were identified using original literature and reference collections by specialist collaborators.

### AHE sequencing and data processing

Cicada specimens were sequenced by Anchored Hybrid Enrichment (AHE; Lemmon et al. 2012) at the Center for Anchored Phylogenomics (www.anchoredphylogeny.com) except for specimen PL623 (*Derotettix mendosensis* Berg), for which sequence reads were obtained from NCBI (SRA: SRS20953264). We prepared Illumina libraries for cicada specimens following Prum et al. (2015) from genomic DNA. Briefly, DNA was fragmented to 125-400 bp with a Covaris ultrasonicator, ligated to Illumina adaptors with 8 bp indexes, and the libraries pooled in groups of ∼24 samples. Each library pool was enriched using the Auchenorrhyncha probe set developed by Dietrich et al. (2017) and described in Simon et al. (2019). Enriched reads were sequenced on an Illumina NovaSeq6000 with a PE150bp protocol. All sequence reads were assembled using the data processing pipeline (https://github.com/markstukel/Simon-target-capture-pipeline) described in Stukel et al. (2024) with a few modifications. To summarize, Illumina adaptors were trimmed from the reads with Trimmomatic v. 0.36, the trimmed reads merged with BBMerge v. 37.41, and the reads from each cicada sample assembled using SPAdes v. 3.12.0 (Bankevich et al. 2012; Bolger et al. 2014; Bushnell et al. 2017). The assemblies were queried for AHE locus sequences using BLASTN, the top matching sequences for each locus in the assemblies extracted using AliBaSeq, and the extracted sequences aligned using MAFFT v. 7.4 (Camacho et al. 2009; Katoh and Standley 2013; Knyshov et al. 2021). Finally, the alignments were clustered using CDHit v. 4.8.1 (Fu et al. 2012) to remove redundant sequences and close in-paralogs. UPhO (Ballesteros and Hormiga 2016) was used to identify more divergent paralog groups, which were later split off into new locus alignments following the decision process outlined by Stukel et al. (2024). The modifications to the pipeline involved the paralogy identification step to accommodate the six pairs of taxa that Owen et al. (2022) identified as having contamination: *Zammara* cf. *erna* and *Platypleura octoguttata*, *Durangona tigrina* and *Hamza ciliaris*, *Quintilia wealei* and *Talcopsaltria olivei*, *Tettigomyia vespiformi*s and *Adusella insignifera*, *Parnisa* sp. and *Kikihia rosea*, and *Lembeja paradoxa* and *H. ciliaris* (see Supplementary Methods for details).

We conducted our final alignment processing and screened the resulting locus alignments for the number of nuclear loci recovered. We used ALiBaSeq and MAFFT to re-extract alignments with 300 bp flanking regions without paralogs or contaminants from the assemblies. Misaligned or misassembled regions in these assemblies were removed using HMMCleaner (Franco et al. 2019) and the ends of the alignments were trimmed until 75% of the taxa were present using a custom script. As several of our assemblies were sequenced from museum specimens, we were concerned about unreliable sequence data from poor-quality DNA. Because taxa yielding abnormally few loci seemed the most likely to be unreliable, we removed taxa with fewer than 100 recovered loci from all alignments (taxa that were removed are not included in the specimen table). Subsequently, we screened the new alignments for new contamination not detected by Owen et al. (2022) using their detection pipeline and removed putative contaminants from the affected loci.

### Assembly of mitochondrial genomes from AHE bycatch

In addition to extracting nuclear AHE loci, we assembled mitochondrial genomes from off-target AHE capture data in the SPAdes assemblies (Simon et al. 2019; Haji et al. 2022; Stukel et al. 2024). Mitochondrial contigs in the assemblies were identified using a *Derotettix mendosensis* reference genome (Łukasik et al. 2019), Reads were mapped to these contigs using BWA (Li and Durbin 2009), and the mapped reads were assembled with SPAdes (Bankevich et al. 2012). The reassembled contigs were aligned to the reference with MAFFT, the contigs merged into a single genome sequence in Geneious, and misassembled regions removed (Kearse et al. 2012). MITObim (Hahn et al. 2013) was then used to extend the contigs with unassembled sequence reads, the extended genome sequences aligned in MAFFT, and the resulting alignment edited manually in Geneious to remove misassembled regions and adjust reading frames to match the reference genome. Mitochondrial genomes from the *Kikihia*, *Maoricicada*, *Rhodopsalta*, and *Derotettix* were obtained from NCBI (GenBank accessions MG737807, OR413956, OR459873, OR459874, OR459906, OR413963) rather than re-extracting them from our assemblies.

### Preliminary phylogenetic analysis

Preliminary phylogenetic analyses were conducted to identify support for various cicada clades for subsequent analyses. We constructed a concatenated maximum-likelihood (ML) phylogeny for the nuclear AHE data with IQ-Tree v. 2.2.2 (Minh et al. 2020). The data were partitioned by gene, and the best model and partitioning scheme was selected using the -*m TESTMERGE* setting in ModelFinder, which uses the PartitionFinder “greedy” algorithm (Chernomor et al. 2016; Kalyaanamoorthy et al. 2017; Lanfear et al. 2017). To save computation time, the relaxed hierarchical clustering algorithm was used to examine only the top 10% of partition merging schemes (Lanfear et al. 2014). AHE loci were not partitioned by codon due to their typically short lengths. Branch support statistics were estimated using 1000 ultrafast bootstrap (UFB) and 1000 Shimodaira-Hasegawa-like approximate likelihood ratio test (SH-aLRT) replicates (Guindon et al. 2010; Hoang et al. 2018). Species trees were estimated from the nuclear AHE data using ASTRAL v. 5.7.3 and SVDQuartets (Chifman and Kubatko 2014; Zhang et al. 2018). For ASTRAL, individual gene trees were first estimated in IQ-Tree with model selection using ModelFinder and 1000 SH-aLRT replicates, and then all gene tree branches with 0% SH-aLRT support were removed with *newick-utilities* to reduce the effect of gene tree estimation error. Based on the recommendations of Simmons and Gatesy (2021), we chose this SH-aLRT threshold rather than a bootstrap support threshold. While majority-rule consensus trees have been argued to best represent uncertainty for a set of trees (Berry and Gascuel 1996; Holder et al. 2008), ASTRAL has been shown to have better performance with ML trees rather than with bootstrap consensus trees (Sayyari and Mirarab 2016). The SVDQuartets analysis was run in PAUP* v. 4.0a (Wilgenbusch and Swofford 2003) using the concatenated alignment and 100 multilocus bootstrap replicates to estimate branch supports. Although methods for estimating branch lengths have recently been developed for SVDQuartets (Peng et al. 2022), we only estimated the topology for our SVDQuartets phylogeny.

A ML mitochondrial phylogeny was constructed from the recovered mitochondrial genomes using IQ-Tree v. 2.2.2. The mitochondrial genomes were partitioned by gene and codon, and the same ModelFinder settings as the concatenated nuclear analysis above were used to merge partitioning schemes. Branch support statistics were again estimated using 1000 UFB and SH-aLRT replicates.

### Bayesian species tree inference and divergence time estimation

Bayesian species tree inference and divergence time estimation were conducted in BEAST v. 2.7.5 (Bouckaert et al. 2019). Clades with > 95% branch support in IQ-Tree and ASTRAL analyses and > 85% branch support in SVDQuartets were used as monophyly constraints for fossil placements. Divergence time estimation was performed in an MSC framework to avoid inflating terminal branch lengths when conducting inference with concatenation (Carruthers et al. 2022). Due to the size of the dataset, a divide and conquer approach was used, in which a reduced taxon set was used to estimate a backbone tree. Taxon sets representing subclades from that backbone were then used to estimate subtrees which were grafted onto the backbone tree. To ensure that the subtrees had dates compatible with the backbone tree, we used approximations of the divergence time posterior distributions in the backbone tree as prior information for the subtrees. This necessitated using different subsets of loci between the backbone tree and the subtrees to avoid “double-dipping” (i.e., using the same data for determining both prior and likelihood). The backbone taxon set consisted of 20 taxa spanning the deepest splits among and within the five cicadid subfamilies (Table S1). The subtree taxon sets consisted of the taxa belonging to Tibicininae, Cicadettinae, Tettigomyiinae, Tacuini, and Cicadinae without Tacuini. For the backbone and subtree taxon sets, species tree inference was performed using StarBeast3 v. 1.1.8 with a relaxed clock, using a fossilized birth-death (FBD) tree prior from Sampled-Ancestors v. 2.1.1 for time calibration and substitution model averaging from bModelTest v. 1.3.3 (Gavryushkina et al. 2014; Bouckaert and Drummond 2017; Douglas et al. 2021, 2022).

Additional details for the different taxon set analyses are explained in the Supplementary Methods.

## Results

### AHE sequencing and processing

Our final dataset recovered 160 taxa and 490 nuclear AHE loci (Table S1). The taxon sampling of Cicadidae from Owen et al. (2022) was expanded from 100 ingroup taxa to 158 ingroup taxa while retaining the two Tettigarctidae outgroup taxa (Table S1). Our 158 ingroup taxa comprise 98 taxa carried over from Owen et al. (2022) and 60 additional taxa (*Magicicada septendecula* and *Lamotialna condamini* specimens from the previous study were removed as we had difficulty processing their AHE sequence reads). The 60 additional species represent one additional subfamily, three additional tribes, and 37 additional genera to the study in Owen et al. (2022). The final taxon sample included all five subfamilies of Cicadidae, 47 of 57 tribes (82%), and 128 of 514 ingroup genera (25%) (World Auchennoryncha Database, accessed 26 March 2026). After removing taxa from loci that were previously identified as contaminated (Owen et al. 2022), we identified two additional pairs of taxa as contaminated. These pairs were *Dorisiana* sp. and *Heteropsaltria aliena* (64 loci) and *Prasinosoma* sp. and *Prasia faticina* (94 loci). Interestingly, the Lemmon Lab AHE sample IDs for each pair are the same numerical distance apart (24 IDs), offset by two (I31409 & I31433 vs I31407 & I31431). The number of loci recovered for each taxon after removing contamination ranged from 94 to 463, with a mean of 396. The taxon occupancy across the nuclear loci ranged from 74 to 159 taxa, with a mean of 129. The length of the nuclear loci ranged from 153 to 1,381 bp, with a mean of 326 bp. The total nuclear alignment length was 159,796 bp. We recovered 114 mitochondrial genomes from AHE bycatch; including the GenBank sequences, this brought our entire mitochondrial alignment to 120 taxa. The total alignment length for the mitochondrial genomes was 14,620 bp.

### Phylogenetic analyses

The ASTRAL, SVDQuartets, and concatenated ML nuclear phylogenies, and the ML mitochondrial genome phylogeny, were all largely congruent (Figs. 1-2). All four of these analyses recovered the same topological arrangement of cicada subfamilies. Other previously-established relationships included deep splits within the subfamilies Cicadettinae and Cicadinae: Cicadettine was divided into two large clades and Tacuini was separated from the rest of Cicadinae. Most disagreements in topology among analyses were relationships that were strongly supported in few or no analyses. These included the relationships of the deep radiation of Australian Cicadinae tribes, the position of *Antankaria*, and the position of *Durangona*. Although IQ-Tree ultrafast bootstraps (UFB), SH-aLRT scores, ASTRAL local posterior probabilities, and regular bootstraps are different measures of branch support that are difficult to compare directly, the SVDQuartets tree displayed much less overall branch support than the ASTRAL or nuclear concatenated ML trees.

**Figure 1:**
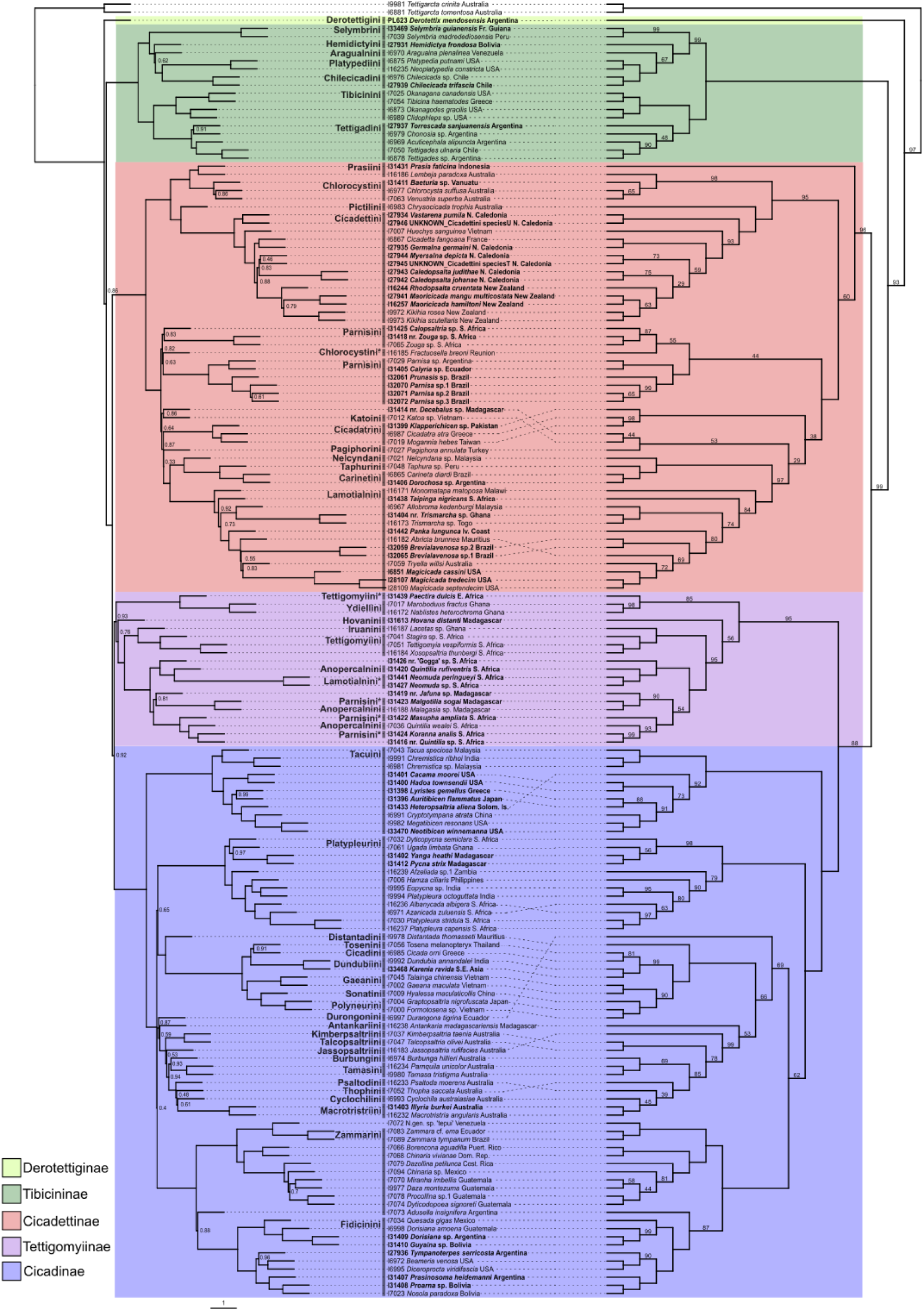
Nuclear ASTRAL and SVDQuartets phylogenies Nuclear ASTRAL and SVDQuartets phylogenies plotted face-to-face. Taxa in bold are newly added relative to Owen et al. (2022). Clade colors indicate subfamily (color key in lower left). Current tribal classifications are labeled; taxa requiring reclassification marked with asterisks. Left: ASTRAL phylogeny constructed from 490 nuclear gene trees. Internal branch lengths are in coalescent units (scale on bottom left). Terminal branch lengths are arbitrary. Nodal support statistics with ASTRAL local posterior probability below 1 are labeled. Right: SVDQuartets cladogram constructed from concatenated alignment of 490 nuclear loci. Branch lengths are arbitrary. Branch support statistics with bootstrap proportion below 100% are labeled.

**Figure 2:**
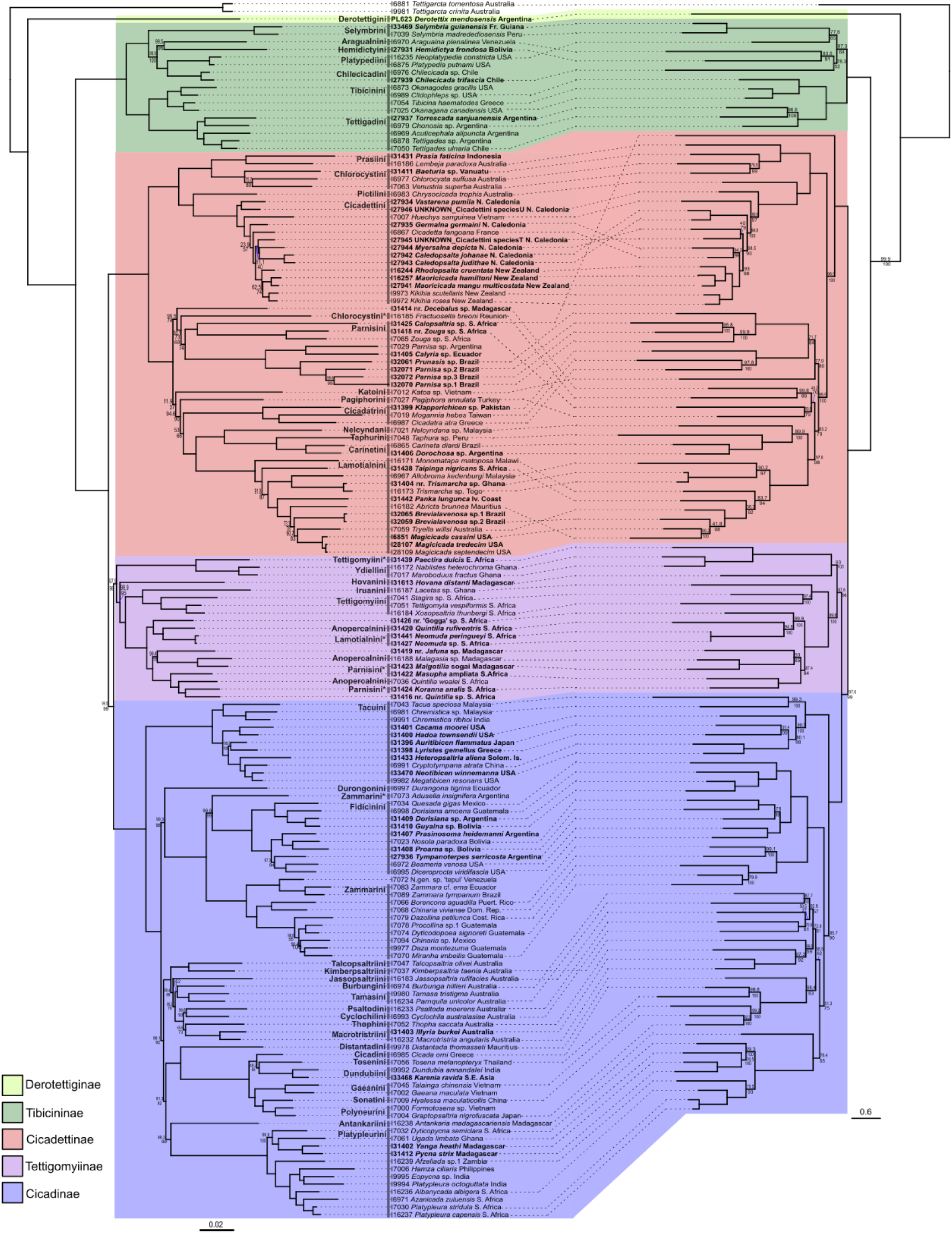
Concatenated nuclear and mitochondrial genome IQ-Tree phylogenies Concatenated nuclear and mitochondrial genome IQ-Tree maximum-likelihood (ML) phylogenies plotted face-to-face. Taxa in bold are newly added relative to Owen et al. (2022). Clade colors indicate subfamily (color key in lower left). Current tribal classifications are labeled; taxa requiring reclassification marked with asterisks. Branch support statistics on both trees are SH-aLRT above, Ultrafast bootstrap support below, with only branches under 100% for either support value labeled. Left: concatenated nuclear IQ-Tree ML phylogeny constructed from 490 nuclear loci. Branch lengths are in substitutions/site (scale on bottom left). Right: IQ-Tree ML phylogeny constructed from mitochondrial genomes obtained from AHE bycatch or NCBI. Branch lengths are in substitutions/site (scale on bottom right). Note that for some taxa, mitochondrial genomes were not recovered.

### Bayesian species tree inference and divergence time estimation

The three versions of the backbone analysis (standard FBD; Skyline-FBD with diversified sampling and two transition points; Skyline-FBD with diversified sampling only) yielded identical summary tree topologies, but slightly different divergence time estimates (Table S4).

The standard FBD analysis yielded slightly older estimated ages for the root and deeper clades, but considerably younger estimated ages for shallower clades. The two Skyline-FBD analyses estimated similar ages for most clades but differed slightly for clades that were close to the Jurassic-Cretaceous and Cretaceous-Paleogene transition points. A closer examination of the analysis with transition points showed unusually shaped posteriors for clade heights very close to the transition points: posterior density was concentrated on one side of the transition and diffused in a long tail on the other side. We believe this is an artifact from the diversification, turnover, and sampling rates changing as they cross those transition points. We therefore selected the clade heights from the Skyline-FBD with diversified sampling only analysis to serve as root or origin age priors for the subtree analyses.

Excluding the branches that were constrained for fossil placement, the combined StarBeast3 tree largely agreed with the ML, ASTRAL, and SVDQuartets phylogenies described above (Fig. 3). Areas that had lower support in the StarBeast3 tree than the previous trees included the sister relationship of Tettigomyiinae + Cicadinae and the monophyly of Tettigomyiinae. However, the posterior probabilities in the StarBeast3 tree should be compared to ML branch support statistics like bootstrap percentages or SH-aLRT scores with caution because, as discussed earlier, branches in the StarBeast backbone or subtree phylogenies were inferred using information from only 50 nuclear genes instead of the full 490 nuclear genes for computational tractability.

**Figure 3:**
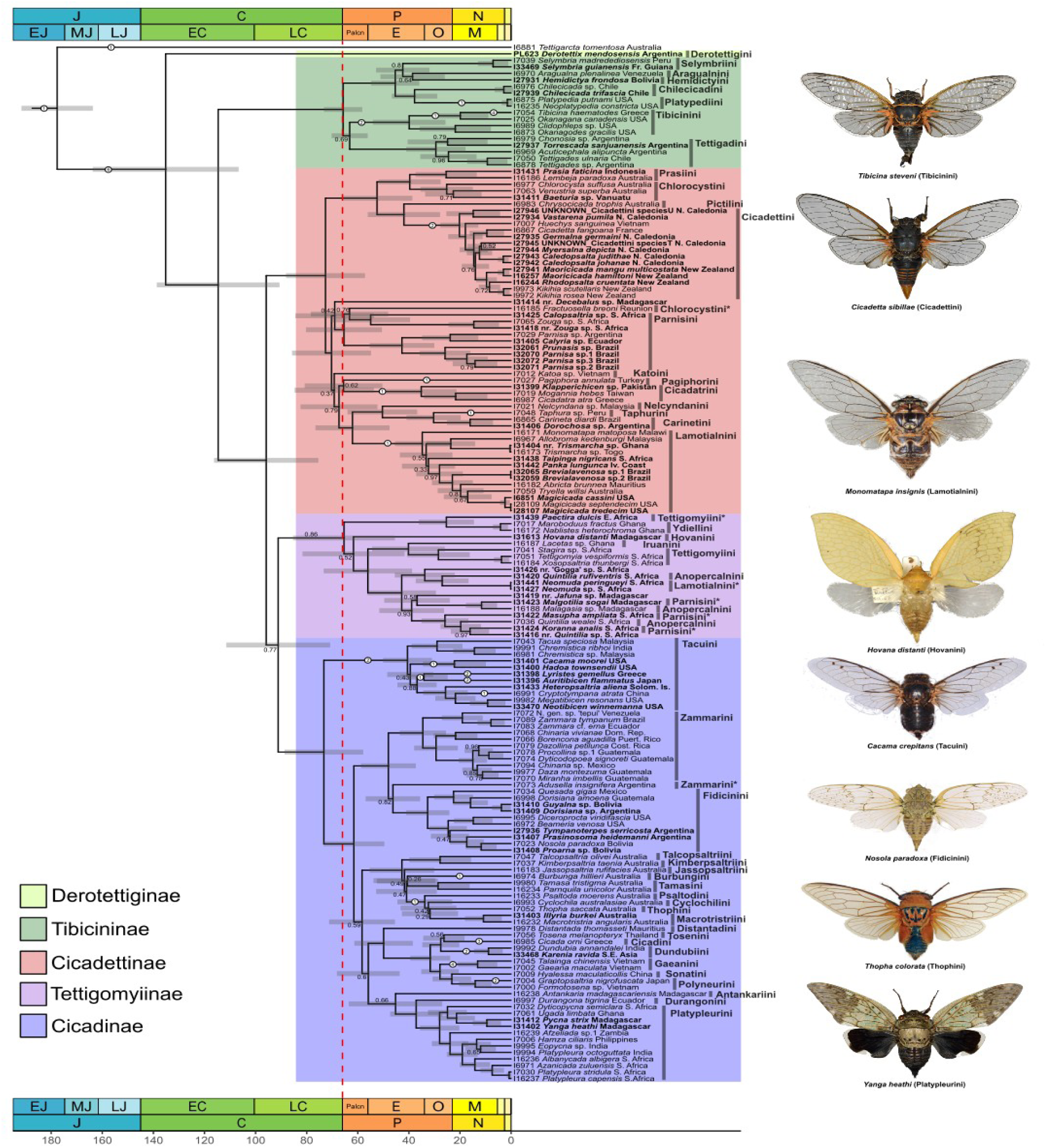
Composite time-calibrated StarBeast3 phylogeny Time calibrated StarBeast3 phylogeny constructed from backbone and subtrees. Taxa in bold are newly added relative to Owen et al. (2022). Clade colors indicate subfamily (color key in lower left). Branches with posterior probabilities below 1 are labeled. Current tribal classifications are labeled; taxa requiring reclassification marked with asterisks. Node heights are common ancestor heights, with 95% HPD intervals displayed as node bars. White circles with numbers indicate branches constrained with fossils and the number of fossils assigned to the branch. Period and epoch data from International Commission on Stratigraphy (v2023/09) plotted at the top and bottom with standard abbreviations (Quaternary, Pliocene, and Pleistocene not labeled). Red dotted line indicates K-Pg transition. Cicada photos highlight exemplar genera and species from major clades in the tree (not to scale). Photo credits: Tibicina, Cicadetta, Monomatapa, Hovana, Yanga: S.P.; Cacama, Nosola, Thopha: D. Marshall.

The divergence times for our combined StarBeast3 tree showed a crown age for Cicadoidea in the early Jurassic (170 Ma, HPD 164.8-192.8 Ma), a crown age for Cicadidae in the early Cretaceous (135.2 Ma, HPD 105.3-162.5 Ma), and crown ages for the four non-monotypic Cicadidae subfamilies at the end of the Cretaceous or at the very start of the Paleocene (Tibicininae: 65.7 Ma, HPD 59-74 Ma; Cicadettinae: 72.6, HPD 57.8-88.7 Ma; Tettigomyiinae: 65.4 Ma, HPD 45.4-85.3 Ma; Cicadinae: 65.4 Ma, HPD 45.4-85.3 Ma). The subfamilies Cicadinae and Cicadettinae, which had estimated crown ages in the Cretaceous, had major radiations estimated to originate at or very close to the K-Pg transition. The crown ages for most of the subtree analyses were consistent with corresponding ages in the backbone tree. As the priors for the crown ages in the subtrees were based on the posterior distributions in the backbone tree, this indicates that the data in these subtrees did not overrule the prior. The only exception to this was the Tettigomyiinae subtree, whose crown age posterior covered only the more recent half of its crown age prior (Table S4).

## Discussion

### Increased phylogenetic resolution of Cicadidae relationships

This study provides increased resolution for relationships within Cicadidae, particularly of the polytomy between Tettigomyiinae, Cicadettinae, and Cicadinae (Marshall et al. 2018; Owen et al. 2022). Using five genes, Marshall et al. (2018), were first to identify Tettigomyiinae as a subfamily, but recovered it as non-monophyletic and in a polytomy with Cicadettinae and Cicadinae. A monophyletic Tettigomyiinae sister to Cicadinettinae + Cicadinae was supported by additional genes (Simon et al. 2019) but with fewer sampled species in comparison with Marshall et al. (2018) Later, phylogenomic analyses of dramatically increased genomic data provided equal support for Cicadettinae + Tettigomyiinae and Cicadinae + Tettigomyiinae (Owen et al. 2022). Our analyses point to Tettigomyiinae + Cicadinae as the resolution of this polytomy. This is likely a result of increased taxon sampling in Tettigomyiinae in this study, which went from eight (Owen et al. 2022) to 19 taxa.

The topologies in our concatenated ML, ASTRAL, and SVDQuartets phylogenies are quite similar, clarifying many relationships among taxa within Cicadidae and identifying areas needing taxonomic revision within the family (Figs). Our results provide molecular confirmation of the placement of *Hovana* and its tribe Hovanini into Tettigomyiinae (Sanborn et al. 2020). Our results also demonstrate that several African genera currently assigned to tribes in Cicadettinae also belong to Tettigomyiinae, increasing the recognized diversity of this subfamily. These include *Neomuda*, currently assigned to Lamotialnini, *Koranna*, currently assigned to Parnisini, *Malgotilia*, currently assigned to Parnisini, and *Masupha*, also currently assigned to Parnisini.

The taxa assigned to Parnisini are especially significant, as Marshall et al. (2018) identified this tribe as having a problematic distribution, with the true members of the tribe probably endemic to South America. Interestingly, Sanborn (2023) removed *Brevialavenosa* from Lamotialnini and placed it in the Taphurini (Cicadettinae) saying, “The transfer of both Brazilian species previously assigned to *Abroma* means there are no members of Lamotialnini remaining in Brazil or South America confirming the tribal distribution as outlined in Marshall et al. (2018).” Our results, however, clearly place *Brevialavenosa* into the Lamotialnini, making it the second genus in this tribe, other than *Magicicada*, to be confirmed in the Americas. During the preparation of this manuscript, Sanborn (2025) moved *Brevialavenosa* back into the Lamotialnini on the basis of our preliminary results and after morphological re-evaluation. One other genus from the Americas, *Chrysolasia*, is currently classified in Lamotialnini (Moulds 2003), but we were unable to extract DNA from the only remaining specimens (loaned by the NHMUK). Our results also agree with Owen et al. (2022) in placing *Aragualna* in Tibicininae and provide phylogenomic support for the placement of *Hemidictya* (Hemidictyini) in Tibicininae by Sanborn et al. (2020). In Cicadinae, the placement of *Antankaria* (Antankariini) and *Durangona* (Durangonini) differ across analyses and receive low branch support in the ASTRAL and SVDQuartets trees (Fig. 1). The concatenated tree shows Antankariini (from Madagascar) as sister to Platypleurini (primarily from Africa and Madagascar) and Durangonini (from South America) as sister to Fidicinini + Zammarini (South and Central America), both with high support, which makes the most sense biogeographically. The other phylogenetic analyses place these two taxa elsewhere near taxa from more distant geographic areas with lower support.

Additionally, we recovered *Adusella insignifera* as a sister-group of Fidicinini and *A. insignifera* + Fidicinini as a sister-group of the remaining Zammarini (Figs. 1-3). Our result suggests that Zammarini including Adusella is non-monophyletic, consistent with the relationship recovered by Goemans (2016) and Owen et al. (2022). Considering the position of *A. insignifera* in the phylogenetic trees (Figs. 1-3) and the morphological differences between species of *Adusella* and the other genera currently classified within Zammarini (Goemans 2016; Sanborn 2019) and Fidicinini, the proposal of a new tribe to classify this genus appears appropriate (Ruschel in prep.).

### Divergence times and the age of Cicadidae

Our results suggest that Cicadidae arose in the early Cretaceous and underwent diversification in the late Cretaceous and early Paleogene, coinciding with the K-Pg mass extinction event. Previous studies have generally assumed an early Paleocene or late Cretaceous origin for Cicadidae based on the family’s fossil evidence and phylogenetic-biogeographic analyses (Moulds 2018; Ruschel and Campos 2019; Simon et al. 2019; Wang et al. 2022). The oldest fossil unambiguously assigned to Cicadidae, *Davispia bearcreekensis*, is from the late Paleocene (Cooper 1941); a late Cretaceous fossil of a first-instar nymph, *Burmacicada protera*, ambiguously represents Cicadidae or Tettigarctidae (Moulds 2018). We estimate that Cicadidae may have an early Cretaceous crown age, which would imply a nearly 70 Myr “phylogenetic fuse” or “ghost lineage”, meaning a time interval between when diversification began and when it first appeared in the fossil record (Cooper and Fortey 1998; Marshall 2019). The proposed existence of long phylogenetic fuses has recently been criticized as an analytical artifact stemming from interaction among too-broad calibration densities (Budd and Mann 2020, 2023). Our methods sought to avoid this artifact by using FBD process to calibrate the tree, rather than assigning nodes subjective calibration densities. Furthermore, *D. bearcreekensis* is assigned to an extant tribe (Tibicinini, Tibicininae) that arose after the split between Derotettiginae and all other Cicadidae subfamilies, which lends more credence to the hypothesis that Cicadidae is considerably older than this fossil.

Our results, likely influenced by our selection of fossil taxa, place the crown age of Cicadoidea in the Early-to-Mid Jurassic. Several fossils assigned to Tettigarctidae have been identified from the Early Jurassic (Shcherbakov 2009), and a few have been identified from the Late Triassic (Lambkin 2019; Jouault et al. 2024). One such fossil, *Sanmai*? *zetavena* (Jouault et al. 2024), was assigned to a genus that morphological phylogenetic analysis suggests is a stem-cicadid (Jiang et al. 2024). While these fossils may push the crown age of Cicadoidea into the Triassic, we elected not to use these early tettigarctid fossils in our analysis for several reasons. Jiang et al. (2024) showed that many morphological characters used to assign fossils to Tettigarctidae are plesiomorphic, and that many fossils currently assigned as such may be stem-cicadoids, stem-tettigarctids, or stem-cicadids. Given that these Triassic and Early Jurassic fossils are quite fragmentary, usually consisting of only complete or partial forewing, we believe their phylogenetic placement is questionable and they may in fact be stem-cicadoids. We therefore elected to include only Mesozoic fossils that had been included in the phylogenetic analysis of Jiang et al. (2024).

Although our use of the tip-dating FBD process avoids artifacts associated with node-dating approaches, there is evidence that the inferred crown age for Cicadidae is slightly inflated. If the inferred crown age of Cicadidae were correct, this would suggest a preservation or discovery bias towards stem-group taxa during the Cretaceous given that there are several stem-group fossils known from the long interval between the Cicadidae crown age and the oldest fossil belonging to Cicadidae. Our youngest fossils from the stem-Cicadidae lineage are also at least 40 Myr younger than their split from the lineage leading to crown Cicadidae (Jiang et al. 2024). However, such long terminal branches for sampled stem lineages might not be entirely problematic, because our included stem-Cicadoidea fossil is 80 Myr younger than the inferred split between Cicadidae and Tettigarctidae, and approximately 60 Myr younger than the oldest stem-fossils assigned to those families.

Our FBD analyses were conducted to minimize any possible artifacts such as deep root attraction, in which violations of model assumptions inflate the ages of deeper nodes (Ronquist et al. 2016; Matschiner 2019; Luo et al. 2023; Zhang et al. 2023). The specific model violations involve the assumption that extant tips are sampled randomly and that the diversification, turnover, and fossil sampling rates of the FBD process are constant through time. In our divergence time analysis strategy, the backbone tree analysis was most likely to be affected by these model violations. To minimize the effect of deep-root attraction, we used the age estimates from the backbone tree analysis that incorporated diversified sampling (Zhang et al. 2016) and changing process rates through time (Zhang et al. 2023). The diversified sampling extension of the FBD model incorporates a cut-off age before the present, after which no speciation or fossil sampling events are allowed to occur (Zhang et al. 2016, 2023). However, we did not incorporate the diversified sampling FBD model in the subtree analyses. The subtree datasets incorporated many very recent fossils and consisted of taxon samples that were intermediate between random and diversified sampling, which would result in the cut-off age being so young as to be irrelevant. As a result of these analytical choices, we believe that our age estimates for Cicadidae may be inflated by deep root attraction, but only modestly so.

Additional precision for cicada divergence times could be obtained with more sophisticated fossil placements. The fossil placements in our tree are entirely determined by the tree prior through monophyly constraints, which themselves are based on taxonomic assignments by specialists. Incorporating morphological data into a combined molecular and morphological analysis would allow more precise fossil placement, and information from the morphological evolution rate could shrink both the stem-lineage branch lengths and the crown age estimate (Gavryushkina et al. 2017; Luo et al. 2020). While a morphological data matrix was available for the stem-group fossils and many extant species in our dataset (Moulds 2005; Simon et al. 2019; Jiang et al. 2024), we lacked morphological data for the crown fossils for Cicadidae used in this analysis. Scoring the same morphological character data for the crown-group fossils to complement the stem-fossils and extant species would allow for future refinement of divergence times for Cicadidae through a combined data analysis and provide exploration of the history of morphological evolution in Cicadidae. Unfortunately, many of the known fossils assigned to Cicadidae are only fragmentary, making this approach problematic.

Even accounting for modest inflation of divergence times due to deep root attraction, Cicadidae clearly have a Cretaceous origin and present-day diversity within the Cicadidae subfamilies originated after the Cretaceous-Paleogene mass extinction. North American placental mammals also experienced a spike in turnover associated with the K-Pg mass extinction (Pires et al. 2018). While not a global mass extinction, the cooling and drying sparked by the establishment of the Antarctic Circumpolar Current and a drop in global CO_2_ levels at the Eocene-Oligocene boundary was associated with high turnover in the Australian pygopodoid gecko radiation (Brennan and Oliver 2017) and initiated a period of exuberant episodic diversification of the Australian desert fauna (e.g., the *Pauropsalta* cicada complex of 11 genera; Owen et al. 2017).

As is the case for birds (Brocklehurst and Field 2024) and placental mammals (Carlisle et al. 2023), frogs (Feng et al. 2017), and snakes (Klein et al. 2021), the stem-lineages for all five Cicadidae subfamilies were extant prior to the mass extinction event and crossed the boundary. Four of the five subfamilies diversified around and after the transition boundary, which raises the question why the fifth subfamily, Derotettiginae, did not proliferate. Simon et al. (2019) propose that during a period of mid-Miocene habitat change, the Derotettiginae faced competition from cicadas in the Tibicininae that were better adapted to the new habitat.

The sister family to Cicadidae, Tettigarctidae, declined after the K-Pg transition. The extensive Mesozoic fossil record of Tettigarctidae indicates that it was a thriving Eurasian lineage prior to the K-Pg mass extinction, while the comparatively few Cenozoic fossils suggest that the lineage was in decline after the transition (Shcherbakov 2009; Moulds 2018). However, some Mesozoic fossils currently assigned to Tettigarctidae may instead represent the stem-lineage of Cicadidae (Jiang et al. 2024). Simulation results suggest that stem groups decline rapidly after the origination of the crown group, and that mass extinctions accelerate this process (Budd and Mann 2020; Hoyal Cuthill et al. 2020). The fossil record of insects suggests that mass extinctions are associated with periods of high turnover of insect lineages rather than absolute reductions in insect diversity (Labandeira 2005; Béthoux 2009; Nicholson et al. 2015; Condamine et al. 2016; Wang et al. 2016; Montagna et al. 2019; Schachat and Labandeira 2021), a phenomenon that is compatible with the results presented here. The Cenozoic aridification of Australia is associated with extinction of mesic-adaptive lineages (Byrne et al. 2011). Since the extant species of Tettigarctidae are restricted to alpine mesic habitat in southeastern Australia and Tasmania, we speculate that these climatic changes may have contributed to the decline of Tettigarctidae.

### Comparison with Cicadidae divergence times estimated through node-dating

Our divergence times for lineages within Cicadidae using the FBD model tip-dating method are largely congruent with previously published studies using node-dating approaches. Our estimated divergence time for Platypleurini (28.0 Ma; HPD: 20.3-35.7 Ma) is very close to previously published ages estimated using node dating and molecular clocks (Price et al. 2019). Similarly, we estimated a divergence time of 40.6 Ma (HPD: 32.0-51.1 Ma) for Tacuini, which is slightly older, but with a considerably narrower range, than the estimate from Hill et al. (2015). The age estimates for divergences within the tribe are also comparable. Our results are also congruent with the previously published divergence times for the radiation of the tribes within Asian Cicadinae (Hill et al. 2021; Wang et al. 2024). We estimate that the tribes of this radiation (Cicadini, Dundubiini, Gaeanini, Polyneurini, Sonatini, and Tosenini) began to diversify approximately 31.7 Ma (HPD: 24.2-39.8 Ma), which is almost identical to the age inferred by Hill et al. (2021). Our age estimates for shallower divergences within this radiation are also consistent with previous estimates (Hill et al. 2021; Wang et al. 2024). The congruence of our age estimates with previous estimates is likely related to the overlap in fossil calibrations used, despite the difference in dating method. Our analysis included two fossils for Polyneurini, one of which was used in a previous node-dating estimate (Wang et al. 2024), and we included twelve fossils across all of the Asian tribes, five of which were used in a previous study (Hill et al. 2021). Our sampling for Cicadettini excluded some clades that are sister to the rest of the tribe, meaning that our Cicadettini sample does not include the deepest splits in the tribe. Our sampling for Cicadettini corresponds to the large clade descending from the Cicadettini radiation (sister to *Chrysocicada*) that does not include the *Pauropsalta* complex (Marshall et al. 2016). We estimated a mean divergence time of 20.2 Ma (95% HPD: 13.1-28.0 Ma) for this clade, which is on the younger end of estimates from the various molecular clock-based analyses of Marshall et al. (2016). Those authors’ estimates suffered from substantial inflation of molecular clock rates, which they tried to rein in using a variety of calibration methods. Our results for Cicadettini have the advantage of the inclusion of a second fossil belonging to the tribe from the Miocene (Moulds et al. 2022), clock rate information from many nuclear loci, the inclusion of many other taxa whose presence can constrain the species tree clock rate in this clade, and a model that accounts for the phylogenetic uncertainty of fossil placement (Gavryushkina et al. 2014; Heath et al. 2014; Zhang et al. 2023).

In a study examining the co-evolution of cicadas and their symbionts, Wang et al. (2022) inferred a crown age for Cicadidae in the late Cretaceous very close to the K-Pg transition (76-66 Ma) based on mitochondrial COI sequences and eight fossil calibrations. This younger date is more consistent with the observed fossil record for Cicadidae, considering the absence of unambiguous crown Cicadidae fossils from the Mesozoic. However, we argue for our older date based on the advantages of the FBD model’s incorporation of fossils into the phylogenetic analysis. Also, Wang et al. (2022) chose to assign *D. bearcreekensis* as a calibration for the root of Tibicininae, despite the fact that this fossil is assigned to an extant tribe within the subfamily.

Furthermore, the maximum age of their tree was constrained by the Misof et al. (2014) estimate of 150 Myr for the age of divergence between Cicadoidea and Cercopoidea. This divergence estimate is certainly an underestimate, as the oldest fossils from Cicadoidea have been dated to over 200 Ma (Shcherbakov 2009; Lambkin 2019; Jouault et al. 2024). More recent time-calibrated phylogenies for Hemipteran insects that incorporate the oldest fossils from Cicadoidea push the divergence between Cicadoidea and Cercopoidea to the late Triassic, around 215 Ma (Johnson et al. 2018). Based on these factors, we believe that the younger date for crown Cicadidae obtained by Wang et al. (2022) is an underestimate.

Osozawa and Wakabayashi (2025) inferred crown ages for Cicadoidea and Cicadidae of 200 Ma and 99 Ma, respectively, using mitochondrial COI and 18S rRNA sequences, twelve geologic calibrations, and thirteen fossil calibrations. However, this phylogeny is not calibrated through typical node-dating approaches, in which calibration densities are constructed around minimum ages determined by the fossil age and maximum ages constrained by prior knowledge or other model parameters (Ho and Phillips 2009). Instead, this phylogeny is a minimum age tree, which avoids the perceived subjectivity surrounding selecting maximum ages for calibration densities and instead calibrates nodes directly to their oldest respective fossils (Klopfstein 2021). Because the minimum age tree does not attempt to estimate the mean or maximum age of a lineage, direct comparison with ages inferred from our FBD analyses or other node-dating studies is difficult. For example, Osozawa and Wakabayashi (2025) estimate the minimum age of Cicadettinae at 30 Ma, which is almost 30 Myr younger than the lower bound of the 95% credible interval in our analyses (Table S4).

Our dating results are congruent with previous node-dating estimates, which raises the question of whether using the FBD model is worth the extra model complexity. We think it is. An advantage of modeling the fossilized birth-death process is that the subjective choices for the node prior calibration densities are replaced with a model incorporating fossil information into the diversification process (Heath et al. 2014). Furthermore, information from all available fossils is incorporated into the estimate rather than simply the oldest fossil assigned to a node.

Our results for Cicadettini and Tacuini show improvement and refinement of age estimates over previously published dates, demonstrating the benefit of using the FBD model.

## Acknowledgements

We thank the numerous collaborators from around the globe who assisted with field work and provided specimens and sequence data for this project, many of whom are listed in the acknowledgements of Owen et al. (2022). We especially thank David Marshall and Kathy Hill, who were invaluable in coordinating specimen collecting efforts. We thank Michelle Kortyna at the Center for Anchored Phylogenomics for assistance with AHE library preparation. We also thank Alexei Drummond and the University of Auckland Centre for Computational Evolution for hosting MS on a Fulbright Fellowship. Computational resources were provided by the University of Connecticut Computational Biology Core. Elizabeth Jockusch, Paul Lewis, and David Marshall provided helpful suggestions and comments on the manuscript. David Marshall and Max Moulds provided comments on the taxonomic summary appendix.

## Data Availability

Data and scripts are available at Dryad Digital Repository:

## Supplementary Information

Table S1: List of specimens

* Taxonomy needing to be updated based on relationships in this study

† Specimen obtained from Natural History Museum, London, UK (NHMUK)

‡ Specimen obtained from Muséum national d’Histoire naturelle (MNHN)

Table S2: List of fossil calibrations

Table S3: Priors for FBD model parameters

Table S4: Differences in divergence times across different StarBeast3 analyses

Online appendix 1: Summary of the subfamilies, tribes and genera including extant and fossil taxa with comments on classification changes since Marshall et al. (2018).

