## Supplementary Methods and Tables for "Phylogenomics and Fossilized Birth-Death Dating Reveals Extensive Post-Cretaceous Worldwide Diversification of Cicadidae (Hemiptera, Auchenorrhyncha)"

*Modifications to AHE sequencing and processing pipeline*

The modifications to the pipeline involved our paralogy identification step and the six pairs of taxa that Owen et al. (2022) identified as having contamination: *Zammara* cf. *erna* and *Platypleura octoguttata*, *Durangona tigrina* and *Hamza ciliaris*, *Quintilia wealei* and *Talcopsaltria olivei*, *Tettigomyia vespiformi*s and *Adusella insignifera*, *Parnisa* sp. and *Kikihia rosea*, and *Lembeja paradoxa* and *H. ciliaris*. During the UPhO step, the contaminated sequences were allowed as in-paralogs, allowing multiple sequences from a contaminated taxon to be included in an ortholog cluster. Each contaminated locus identified by Owen et al. (2022) was examined for the presence of multiple sequences from a taxon in a contaminated pair. If one taxon in a pair had two sequences for a locus, and one of the sequences was identical to the sequence from the other taxon in the pair, we deleted that duplicate sequence. This allowed us to increase our sequence data for these taxa. For contaminated loci that did not have recovered sequences that could be easily identified as contaminants in this way, we followed Owen et al. (2022) and removed the taxon pair from the contaminated locus. These modifications are included in a replacement UPhO.sh script uploaded to Dryad.

*Bayesian species tree inference and divergence time estimation*

For the backbone tree, nuclear AHE loci with all 20 taxa were identified and from them the 50 loci with the highest number of parsimony-informative sites as calculated by PhyKIT (Steenwyk et al. 2021) were selected. Because of the selective taxon sampling and deep splits in the backbone taxon set, we were concerned that the assumption of constant rates through time might result in “deep root attraction” (Ronquist et al. 2016; Matschiner 2019; Luo et al. 2023; Zhang et al. 2023). To address this, three versions of the backbone analysis were compared: an analysis using the standard FBD model with constant rates; a Skyline-FBD model with diversified sampling assumptions and two rate transition points at the Jurassic-Cretaceous and Cretaceous-Paleogene boundaries; and a Skyline-FBD model with diversified sampling but no rate transition points (Stadler et al. 2013; Stadler and Smrckova 2016; Zhang et al. 2016). We included 15 fossil tips for all three versions of the backbone analysis (Table S2, first 15 lines). Eight fossil tips were stem-group taxa identified in a recent morphological phylogenetic analysis (Jiang et al. 2024), consisting of one stem cicadoid, one stem tettigarctid, and six stem cicadids. Seven further tips were fossils of two Cicadinae, two Cicadettinae, and three Tibicininae, all of which represent the oldest unambiguous crown Cicadidae fossils older than our diversified sampling cutoff of 25 Ma (Moulds 2018). To account for stratigraphic age uncertainty, we specified a log-normal prior for each fossil tip such that the published fossil age range spanned the central 95% of the prior distribution (Barido-Sottani et al. 2019). Based on the estimate of 220 Ma for the divergence between Cicadoidea and Cercopoidea in Johnson et al. (2018) we used a log-normal prior for the origin with a mean in real space of 54 and an offset of 166 (the age of the oldest fossil in our dataset). We set *ρ* (extant sampling proportion) to 0.004 based on an estimate of approximately 4,000 extant species of cicadas. We chose uniform priors for *d* (net diversification rate) and *r* (turnover) (Table S3). Due to the paucity of fossils, the fossil sampling proportion *s* was expected to be small, so the number of sampled fossils was divided by 4,000 extant species of cicadas to get a rough approximation for the sampling proportion*,* and a beta distribution with a mean of this value was chosen to be the prior for *s.*

The StarBeast3 analyses were set up similarly for the individual subtree taxon sets. The 50 loci used in the backbone tree were excluded from consideration and the locus selection process repeated for each taxon subset. The nuclear AHE loci containing all taxa in each subset were identified. If there were fewer than 50 loci that contained all taxa for a set, the taxon occupancy threshold was relaxed down to all loci with at least *N*-2 taxa, where *N* is the number of taxa in the subtree taxon set. The top 50 loci with the largest number of parsimony-informative sites were selected from these high-occupancy loci. Because the subtree taxon sets were not as thinly sampled as the backbone taxon set, all analyses were performed using the standard, constant-rates FBD model. We used a total of 36 fossil tips split among the Cicadettinae, Tibicininae, Tacuini, and Cicadinae without Tacuini datasets; no fossil Tettigomyiinae were included as no fossils from this subfamily have yet been identified (Table S2). We accounted for stratigraphic age uncertainty in all fossils in the same way as for the backbone tree. We chose the backbone tree estimated using the diversified sampling Skyline FBD model with no transition points as the source for calibrating the subtrees. The root age or origin age priors for all subtree analyses were set to approximate the posterior ages for the corresponding clades in this backbone tree, with the appropriate offsets based on the oldest fossils (Table S3). Root ages were approximated for clades which we were confident had their deepest split captured in the backbone tree. For clades that we were unsure had their deepest split captured in the backbone tree, the age distribution of the next-oldest split in the backbone tree was approximated as an origin age prior. A lognormal approximation of the backbone Species Tree Relaxed Clock rate posterior was used as the prior distribution for this parameter in each of the subtrees. Values for *ρ* (extant sampling proportion) were set based on the total number of described species in each subtree clade (Catalog of Life, accessed 13 May 2024) rescaled to an estimate of 4,000 total extant species for the whole family (Table S3). Uniform priors for *d* (net diversification rate) and *r* (turnover) were used as in the backbone analysis (Table S3). The fossil sampling proportion *s* was again approximated by dividing the number of sampled fossils in a dataset by the estimated extant species in the clade, and a beta distribution with a mean of this value was chosen to be the prior (Table S3). Lacking Tettigomyiinae fossils, analysis for this subfamily was attempted using a Yule model instead of an FBD model, but preliminary results were unsatisfactory, likely due to our sampling being insufficiently dense. Instead, an FBD model was used with *s* set to zero, which is equivalent to a birth-death model.

All analyses, whether backbone or subtree, were performed with two identical runs each for up to 400 million generations. Because the Tettigomyiinae and Cicadinae without Tacuini datasets became stuck in local optima, three heated chains and one cold chain were used with the CoupledMCMC v. 1.3.1 package with the default swapping value for those datasets to improve convergence (Müller and Bouckaert 2020); all other analyses lacked heated chains. Convergence was determined by comparing the two independent runs for each analysis in Tracer v. 1.7.2 and the clade comparison tool in DensiTree (Bouckaert and Heled 2014). The first 10% of tree samples were removed as burn-in from each run and the species tree files were combined in LogCombiner v. 2.7.5. Fossil tips were removed from each combined species tree files and MCC consensus trees were generated with common ancestor node heights in TreeAnnotator (Heled and Bouckaert 2013). The subtree consensus trees and their associated metadata were grafted onto the backbone tree in R. Because we considered the root age estimates and highest posterior density (HPD) distributions from the different subtrees to be more refined than those from the backbone tree, subtrees were attached to the backbone at their inferred root node height and the corresponding node on the backbone tree was removed. However, the posterior probabilities from the backbone tree for these nodes were kept because we believed they were closer to the true level of uncertainty for these relationships (the posterior probabilities for these nodes in the subtrees were all 1 due to being their roots).

**Supplementary Tables**

**Table S1: List of specimens** (separate file)

* Taxonomy needing to be updated based on relationships in this study

† Specimen obtained from Natural History Museum, London, UK (NHMUK)

‡ Specimen obtained from Muséum national d’Histoire naturelle (MNHN)

**Table S2: List of fossil calibrations**

Because phylogenetic constraints are nested, only the narrowest constraint for each analysis is listed for each fossil. All fossils are necessarily included in constraints for deeper clades that contain the listed constraint.

| **Fossil** | **Age (Ma)**  **Min Max** | | **Phylogenetic constraint** | **Notes** | **Reference** |
| --- | --- | --- | --- | --- | --- |
| *Macrotettigarcta obesa* | 157 | 166 | Backbone: stem Cicadidae |  | Jiang et al. 2024 |
| *Sanmai mengi* | 157 | 166 | Backbone: stem Cicadidae |  | Jiang et al. 2024 |
| *Shuraboprosbole daohugouensis* | 157 | 166 | Backbone: stem Cicadidae |  | Jiang et al. 2024 |
| *Tianyuprosbole zhengi* | 157 | 166 | Backbone: stem Tettigarctidae |  | Jiang et al. 2024 |
| *Cretotettigarcta problematica* | 98.2 | 99.4 | Backbone: stem Cicadidae; in a clade with *Pranwanna*, *Vetuprosbole*, and crown Cicadidae | Placement informed by well-supported published morphological analysis | Jiang et al. 2024 |
| *Eunotalia emeryi* | 98.2 | 99.4 | Backbone: stem Cicadoidea |  | Jiang et al. 2024 |
| *Pranwanna xiai* | 98.2 | 99.4 | Backbone: stem Cicadidae; sister to crown Cicadidae | Placement informed by well-supported published morphological analysis | Jiang et al. 2024 |
| *Vetuprosbole parallelica* | 98.2 | 99.4 | Backbone: stem Cicadidae; in a clade with *Cretotettigarcta*, *Pranwanna*, and crown Cicadidae | Placement informed by well-supported published morphological analysis | Jiang et al. 2024 |
| *Davispia bearcreekensis* | 56 | 59.2 | Backbone and Tibicininae: Tibicinini |  | Moulds 2018 |
| *Lyristes* sp. | 30.2 | 30.5 | Backbone: Tacuini; in a clade with *Hadoa* and *Megatibicen*.  Tacuini: in a clade with *Auritibicen* and *Lyristes* | Placement in backbone tree is the branch of Tacuini where the genus belongs.  Placement in Tacuini tree based on old age of fossil vs. low divergence between extant *Auritibicen* and *Lyristes* | Moulds 2018 |
| *Hadoa grandiosa* | 27.8 | 33.9 | Backbone: Tacuini; in a clade with *Lyristes* and *Megatibicen.*  Tacuini: sister to extant *Cacama* and *Hadoa* | Placement in backbone tree is the branch of Tacuini where the genus belongs. Placement in Tacuini tree is based on old age of fossil vs. low divergence between extant *Cacama* and *Hadoa*. | Moulds 2018 |
| *Lithocicada perita* | 27.8 | 33.9 | Backbone and Tibicininae: Tibicinini |  | Moulds 2018 |
| *Paracicadetta oligocenica* | 27.8 | 33.9 | Backbone: in a clade with *Cicadatra* and *Magicicada.*  Cicadettinae: Pagiphorini | Placement on the backbone tree is the branch of Cicadettinae where the genus belongs. | Moulds 2018 |
| *Platypedia primigenia* | 27.8 | 33.9 | Backbone and Tibicininae: Platypediini | Placement in the Tibicininae tree is based on old age of fossil vs. low divergence between extant *Neoplatypedia* and *Platypedia* | Moulds 2018 |
| *Cicadatra? serresi* | 26.0 | 27.8 | Backbone and Cicadettinae: Cicadatrini | Placement chosen despite tentative assignment of fossil to *Cicadatra* | Moulds 2018 |
| *Tibicina sakalai* | 17.8 | 17.9 | Tibicininae: in a clade with *Okanagana* and *Tibicina* | Placement is based on old age of fossil vs. low divergence between extant *Okanagana* and *Tibicina* | Moulds 2018 |
| *Meimuna protopalifera* | 16 | 23 | Cicadinae (without Tacuini): Dundubiini |  | Moulds 2018 |
| *Tymocicada gorbunovi* | 16 | 23 | Tacuini: Tacuini |  | Moulds 2018 |
| *Paleopsalta ungeri* | 16 | 20.4 | Cicadettinae: Cicadettini |  | Moulds 2018 |
| *Minyscapheus dominicanus* | 15 | 20 | Cicadettinae: Taphurini |  | Moulds 2018 |
| *Cryptotympana incasa* | 11.6 | 16 | Tacuini: *Cryptotympana* |  | Moulds 2018 |
| *Tanyocicada lapidescens* | 11.6 | 16 | Cicadinae (without Tacuini): in a clade with extant Gaeanini and extinct Leptopsaltriini | Placement is based on the close relationship of Leptopsaltriini (the tribe to which the fossil is assigned) and Gaeanini found in Hill & Marshall et al. (2021) | Moulds 2020 |
| *Camuracicada aichhorni* | 11.6 | 13.8 | Tacuini: Tacuini |  | Moulds 2018 |
| *Burbungoides gulgongensis* | 11 | 16 | Cicadinae (without Tacuini): Burbungini |  | Moulds et al. 2022 |
| *Laopsaltria ferruginosa* | 11 | 16 | Cicadettinae: Cicadettini |  | Moulds et al. 2022 |
| *Tithopsaltria titan* | 11 | 16 | Cicadinae (without Tacuini): in a clade with extant Cyclochilini, Macrotristriini, Psaltodini, and Thophini | Placement is based on the close relationship of Arenopsaltriini with these tribes in Marshall et al. (2018) | Moulds et al. 2022 |
| *Lyristes renei* | 8 | 8.5 | Tacuini: *Lyristes* |  | Moulds 2018 |
| *Miocenoprasia grasseti* | 8 | 8.5 | Cicadettinae: Lamotialnini |  | Moulds 2018 |
| *Tibicina gigantea* | 8 | 8.5 | Tibicininae: *Tibicina* |  | Moulds 2018 |
| *Auritibicen* sp. aff. *japonicus* | 5.3 | 11.6 | Tacuini: *Auritibicen* |  | Moulds 2018 |
| *Graptopsaltria inaba* | 5.3 | 11.6 | Cicadinae (without Tacuini): *Graptopsaltria* |  | Moulds 2018 |
| *Yezoterpnosia* sp. aff. *vacua* | 5.3 | 11.6 | Cicadinae (without Tacuini): in a clade with extant Gaeanini and extinct Leptopsaltriini | Placement is based on the close relationship of Leptopsaltriini (the tribe to which the fossil is assigned) and Gaeanini found in Hill & Marshall et al. (2021) | Moulds 2018 |
| *Lyristes? emathion* | 5.3 | 7.2 | Tacuini: *Lyristes* |  | Moulds 2018 |
| *Cicada* sp. aff. *lodosi* | 2.6 | 3.6 | Cicadinae (without Tacuini): *Cicada* |  | Moulds et al. 2023 |
| *Cicada* sp. aff. *orni* | 2.6 | 3.6 | Cicadinae (without Tacuini): *Cicada* |  | Moulds 2018 |
| *Cicada tithonus* | 2.6 | 3.6 | Cicadinae (without Tacuini): *Cicada* |  | Moulds et al. 2023 |
| *Meimuna* sp. | 2.6 | 3.6 | Cicadinae (without Tacuini): Dundubiini |  | Moulds 2018 |
| *Tibicina boulardi* | 2.6 | 3.6 | Tibicininae: *Tibicina* |  | Moulds et al. 2023 |
| *Tibicina* sp. aff. *haematodes* | 2.6 | 3.6 | Tibicininae: *Tibicina* |  | Moulds 2018 |
| *Tibicina lata* | 2.6 | 3.6 | Tibicininae: *Tibicina* |  | Moulds et al. 2023 |
| *Tanna?* sp. | 1.5 | 1.7 | Cicadinae (without Tacuini): in a clade with extant Gaeanini and extinct Leptopsaltriini | Placement is based on the close relationship of Leptopsaltriini (the tribe to which the fossil is assigned) and Gaeanini found in Hill & Marshall et al. (2021) | Moulds 2018 |
| *Graptopsaltria* sp. aff. *nigrofuscata* | 0.5 | 1.1 | Cicadinae (without Tacuini): *Graptopsaltria* |  | Moulds 2018 |
| *Auritibicen bihamatus* | 0.033 | 0.038 | Tacuini: *Auritibicen* |  | Moulds 2018 |
| *Yezoterpnosia nigricosta* | 0.033 | 0.038 | Cicadinae (without Tacuini): in a clade with extant Gaeanini and extinct Leptopsaltriini | Placement is based on the close relationship of Leptopsaltriini (the tribe to which the fossil is assigned) and Gaeanini found in Hill & Marshall et al. (2021) | Moulds 2018 |

**Table S3: Priors for FBD model parameters**

All lognormal distributions are reported with the mean in linear space instead of log space.

| **Analysis** | **Origin** | **Root** | ***ρ* (extant sampling prop.)** | ***d* (diversification rate)** | ***s* (sampling prop.)** | ***r* (turnover)** |
| --- | --- | --- | --- | --- | --- | --- |
| Backbone (all three analyses) | LogNorm(54, 1), offset=166 | NA | 0.005 | Uniform(inf) | Beta(1, 250) | Uniform(1, 0) |
| Cicadettinae | NA | LogNorm(34.1, 0.25), offset=33.9 | 0.04 | Uniform(inf) | Beta(1, 250) | Uniform(1, 0) |
| Cicadinae (no Tacuini) | NA | LogNorm(40, 0.198), offset=23 | 0.03 | Uniform(inf) | Beta(1, 120) | Uniform(1, 0) |
| Tacuini | LogNorm(45, 0.17), offset=28 | NA | 0.04 | Uniform(inf) | Beta(1, 32) | Uniform(1, 0) |
| Tettigomyiinae | LogNorm(90, 0.1) | NA | 0.1 | Uniform(inf) | 0 | Uniform(1, 0) |
| Tibicininae | NA | LogNorm(9, 0.5), offset=56 | 0.06 | Uniform(inf) | Beta(1, 35) | Uniform(1, 0) |

**Table S4: Differences in divergence times across different StarBeast3 analyses**

Common ancestor (CA) heights and highest posterior densities (HPD) for assorted deep clades obtained for each StarBeast3 analysis. The posteriors for the backbone analysis with diversified sampling only (second line of table) were approximated as root or origin prior distributions for the subtree analyses (see Table 2 above).

| **Analysis** | **Cicadoidea CA height (95% HPD)** | **Cicadidae CA height (95% HPD)** | **Cicadettinae CA height (95% HPD)** | **Cicadinae (no Tacuini) CA height (95% HPD)** | **Tacuini CA height (95% HPD)** | **Tettigomyiinae CA height (95% HPD)** | **Tibicininae CA height (95% HPD)** |
| --- | --- | --- | --- | --- | --- | --- | --- |
| Backbone, constant FBD | 197.5 (166.8-247.2) Ma | 169.1 (130.8-221.1) Ma | 92.9 (65.2-125.8) Ma | 83.0 (59.0-111.9) Ma | 45.4 (32.7-61.0) Ma | 109.3 (68.1-154.9) Ma | 71.9 (57.4-92.3) Ma |
| Backbone, Skyline-FBD diversified sampling | 177.7 (164.8-192.8) Ma | 135.2 (105.3-162.5) Ma | 67.7 (49.3-86.2) Ma | 63.2 (40.1-75.4) Ma | 38.2 (31.4-46.8) Ma | 80.2 (45.6, 109.8) Ma | 65.0 (57.0-75.3) Ma |
| Backbone, Skyline-FBD diversified sampling, 2 transition points | 170.0 (161.3-180.2) Ma | 145.2 (123.3-162.3) Ma | 61.2 (45.4-80.4) Ma | 57.7 (47.2-66.0) Ma | 37.8 (25.4-43.7) Ma | 67.4 (27.6-104.8) Ma | 62.0 (57.9-66.0) Ma |
| Subtree, constant FBD | — | — | 72.6 (57.8-88.7) Ma | 61.5 (49.6-73.4) Ma | 40.6 (32.0-51.1) Ma | 65.4 (45.4-85.3) Ma | 65.7 (59.0-74.0) Ma |
