## Supplementary material for "Phylogenomics and Fossilized Birth-Death Dating Reveals Extensive Post-Cretaceous Worldwide Diversification of Cicadidae (Hemiptera, Auchenorrhyncha)": Online appendix 1

**Online Appendix 1: Summary of the subfamilies, tribes and genera including extant and fossil taxa with comments on classification changes since Marshall et al. (2018b).**

**Cicadidae Batsch**

**Comments:** A recent Official Correction was made for the authorship of the family-group names based on *Cicada* as Batsch, 1789 (Official Correction 138 2023).

**Cicadettinae Buckton**

**Carinetini Distant**

*Ahomana* Distant; *Carineta* Amyot & Audinet-Serville; *Dorachosa* Distant; *Guaranisaria* Distant; *Novemcella* Goding; *Toulgoetalna* Boulard.

**Comments:** *Herrera* Distant (Carinetini) was shown to be a junior synonym of *Dorachosa* (Taphurini) and *Dorachosa* was transferred from Taphurini to Carinetini (Kratzer 2024; Sanborn 2025a). *Paranistria* was placed in Parnisini (Sanborn 2024a).

**Chlorocystini Distant**

*Aedeastria* Boer; *Akamba* Distant; *Baeturia* Stål; *Chlorocysta* Westwood; *Cystopsaltria* Goding & Froggatt; *Cystosoma* Westwood; *Decebalus* Distant; *Dinarobia* Mamet; *Euthemopsaltria* Moulds; *Fractuosella* Boulard; *Glaucopsaltria* Goding & Froggatt; *Guineapsaltria* Boer; *Gymnotympana* Stål; *Kumanga* Distant; *Mirabilopsaltria* Boer; *Musoda* Karsch; *Owra* Ashton; *Papuapsaltria* Boer; *Scottotympana* Boer; *Thaumastopsaltria* Kirkaldy; *Venustria* Goding & Froggatt.

**Comments:** *Cephalalna* was transferred to Malagasiini (now Anopercalnini) (Sanborn 2021a). *Conibosa* and *Muda* were transferred to Parnisini and Katoini, respectively (Sanborn 2025a, 2025b).

**Cicadatrini Distant**

*Bijaurana* Distant; *Chloropsalta* Haupt; *Cicadatra* Kolenati; *Emathia* Stål; *Klapperichicen* Dlabola; *Mogannia* Amyot & Audinet-Serville; *Psalmocharias* Kirkaldy; *Shaoshia* Wei, Ahmed & Rizvi; *Taungia* Ollenbach; *Triglena* Fieber; *Vagitanus* Distant.

**Comments:** *Pachypsaltria* is removed from Cicadatrini to Carinetini (Cicadettinae) (Kratzer, 2024). However, Sanborn (2025) provided morphological evidence to classify the genus in Zammarini (Cicadinae).

**Cicadettini Buckton**

*Adelia* Moulds; *Aestuansella* Boulard; *Afromelampsalta* Sanborn & Villet; *Amphipsalta* Fleming; *Atrapsalta* Owen & Moulds; *Auscala* Moulds; *Austropunia* Moulds & Marshall; *Auta* Distant; *Berberigetta* Costa, Nunes, Marabuto, Mendes & Simões; *Birrima* Distant; *Bispinalta* Delorme; *Brevia* Moulds & Marshall; *Buyisa* Distant; *Caledopsalta* Delorme; *Caliginopsalta* Ewart; *Calipsalta* Moulds & Marshall; *Chelapsalta* Moulds; *Cicadetta* Kolenati; *Cicadettana* Marshall & Hill; *Clinata* Moulds; *Clinopsalta* Moulds; *Cognadanga* Moulds & Marshall; *Crotopsalta* Ewart; *Curvicicada* Chou & Lu; *Diemeniana* Distant; *Dimissalna* Boulard; *Dipsopsalta* Moulds; *Drymopsalta* Ewart; *Erempsalta* Moulds; *Euboeana* Gogala, Trilar & Drosopoulos; *Euryphara* Horváth; *Ewartia* Moulds; *Falcatpsalta* Owen & Moulds; *Fijipsalta* Duffels; *Froggattoides* Distant; *Gagatopsalta* Ewart; *Galanga* Moulds; *Gelidea* Moulds; *Germalna* Delorme; *Ggomapsalta* Lee; *Graminitigrina* Ewart & Marques; *Graptotettix* Stål; *Gudanga* Distant; *Haemopsalta* Owen & Moulds; *Hea* Distant; *Heliopsalta* Moulds; *Heremusina* Ewart; *Hilaphura* Webb; *Huechys* Amyot & Audinet-Serville; *Ingcainyenzane* Sanborn & Villet; *Kalarko* Moulds & Marshall; *Kanakia* Distant; *Kikihia* Dugdale; *Kobonga* Distant; *Kosemia* Matsumura; †*Laopsaltria* Moulds, Frese & McCurry; *Limnopsalta* Moulds; *Linguacicada* Chou & Lu; *Maoricicada* Dugdale; *Marteena* Moulds; *Melampsalta* Kolenati; *Melanesiana* Delorme; *Mouia* Distant; *Mugadina* Moulds; *Murmurillana* Delorme; *Myersalna* Boulard; *Myopsalta* Moulds; *Nanopsalta* Moulds; *Neopunia* Moulds; *Nigripsaltria* Boer; *Noongara* Moulds; *Notopsalta* Dugdale; *Oligoglena* Horvath; *Pakidetta* Sanborn & Ahmed; *Palapsalta* Moulds; †*Paleopsalta* Moulds; *Panialna* Delorme; *Paraclinata* Moulds & Marshall; *Paradina* Moulds; *Parvittya* Distant; *Parvopsalta* Moulds & Marshall; *Paulaudalna* Delorme; *Pauropsalta* Goding & Froggatt; *Pedana* Moulds & Marshall; *Pegapsaltria* Moulds & Marshall; *Pericallea* Moulds, Marshall & Hutchinson; *Philipsalta* Lee, Marshall & Hill; *Physeema* Moulds; *Pinheya* Dlabola; *Pipilopsalta* Ewart; *Platypsalta* Moulds; *Plerapsalta* Moulds; *Popplepsalta* Owen & Moulds; *Poviliana* Boulard; *Pseudokanakia* Delorme; *Pseudotettigetta* Puissant; *Punia* Moulds; *Pyropsalta* Moulds; *Relictapsalta* Owen & Moulds; *Rhodopsalta* Dugdale; *Rouxalna* Boulard; *Samaecicada* Popple & Emery; *Saticula* Stål; *Scieroptera* Stål; *Scolopita* Chou & Lei; *Simona* Moulds; *Stellenboschia* Distant; *Strepuntalna* Delorme; *Sylphoides* Moulds; *Takapsalta* Matsumura; *Taurella* Moulds; *Telmapsalta* Moulds; *Terepsalta* Moulds; *Tettigetta* Kolenati; *Tettigettacula* Puissant; *Tettigettalna* Puissant; *Tettigettula* Puissant; *Tibeta* Lei & Chou; *Toxala* Moulds; *Toxopeusella* Schmidt; *Tympanistalna* Boulard; *Ueana* Distant; *Urabunana* Distant; *Uradolichos* Moulds; *Vastarena* Delorme; *Xeropsalta* Ewart; *Xossarella* Boulard; *Yoyetta* Moulds.

**Comments:** Fourteen new genera are proposed to the tribe: *Afromelampsalta* Sanborn & Villet, 2020a; *Austropunia* Moulds &Marshall, 2025; *Brevia* Moulds & Marshall, 2025; *Calipsalta* Moulds & Marshall, 2022; *Cognadanga* Moulds & Marshall, 2025; *Heremusina* Ewart, 2018; *Ingcainyenzane* Sanborn & Villet, 2020b; *Kalarko* Moulds & Marshall, 2022; *Paraclinata* Moulds & Marshall, 2025; *Parvopsalta* Moulds & Marshall, 2022; *Pedana* Moulds & Marshall, 2022; *Pegapsaltria* Moulds & Marshall, 2022; *Pericallea* Moulds, Marshall & Hutchinson, 2022; *Xeropsalta* Ewart, 2018. *Hea* was transferred from Tacuini (= Cryptotympanini) (Cicadinae) to Cicadettini (Cicadettinae) based on morphological evidence (Sanborn, 2023). The new fossils genera, *Paleopsalta* Moulds, 2018 and *Laopsaltria* Moulds, Frese & McCurry 2022 were proposed to the tribe.

**Katoini Moulds & Marshall**

*Katoa* Ouchi; *Muda* Distant.

**Comments:** *Muda* was transferred from the Chlorocystini to the Katoini based on morphological evidence (Sanborn, 2025b).

**Lamotialnini Boulard**

*Abricta* Stål; *Abroma* Stål; *Aleeta* Moulds; *Allobroma* Duffels; *Brevialavenosa* Sanborn; *Chalumalna* Boulard; *Chrysolasia* Moulds; *Hylora* Boulard; *Lamotialna* Boulard; *Lemuriana* Distant; *Magicicada* Davis; †*Miocenoprasia* Boulard & Riou; *Monomatapa* Distant; *Musimoia* China; *Neomuda* Distant; *Oudeboschia* Distant; *Panka* Distant; *Sundabroma* Duffels; *Trismarcha* Karsch; *Tryella* Moulds; *Unduncus* Duffels; *Viettealna* Boulard.

**Comments:** The fossil genus *Miocenoprasia* was transferred from Prasiini to Lamotialnini (Moulds, 2018). The new genus, *Brevialavenosa*, was proposed to Taphurini (Sanborn, 2021b), and later was transferred, as well as *Chalumalna*, from Taphurini to Lamotialnini (Sanborn, 2025c).

**Nelcyndanini Moulds & Marshall**

*Nelbroma* Sanborn; *Nelcyndana* Distant.

**Comments:** The new genus, *Nelbroma*, was proposed to the tribe (Sanborn, 2021a).

**Pagiphorini Moulds & Marshall**

*Pagiphora* Horváth; †*Paracicadetta* Boulard & Nel.

**Comment:** Upon further examination, the fossil genus *Paracicadetta* was considered morphologically close to the extant genus *Pagiphora* (Pagiphorini) instead to *Cicadetta* (Cicadettini) (Moulds, 2018).

**Parnisini Distant**

*Abagazara* Distant; *Acyroneura* Torres; *Adeniana* Distant; *Arcystasia* Distant; *Calopsaltria* Stål; *Calyria* Stål; *Conibosa* Distant; *Crassisternalna* Boulard; *Henicotettix* Stål; *Jafuna* Distant; *Kageralna* Boulard; *Koranna* Distant; *Luangwana* Distant; *Lycurgus* China; *Malgotilia* Boulard; *Mapondera* Distant; *Masupha* Distant; *Paranistria* Metcalf; *Parnisa* Stål; *Prunasis* Stål; *Psilotympana* Stål; *Rhinopsalta* Melichar; *Taipinga* Distant; *Timbaltransversa* Sanborn; *Zouga* Distant.

**Comments:** Three new tribal affiliations: *Paranistria* was transferred from Carinetini to Parnisini (Sanborn, 2024a), *Conibosa* from Chlorocystini to Parnisini (Kratzer 2024; Sanborn, 2025a) and a new genus, *Timbaltransversa*, was proposed to Parnisini (Sanborn, 2021a). *Derotettix* was removed from Parnisini to compose the monogeneric subfamily Derotettiginae (Simon et al., 2019).

**Pictilini Moulds & Hill, 2018**

*Amica* Moulds & Marshall; *Chrysocicada* Boulard; *Pictila* Moulds.

**Comments:** The new genus, *Amica*, was proposed to the tribe (Moulds & Marshall 2025).

**Prasiini Matsumura**

*Arfaka* Distant; *Jacatra* Distant; *Lembeja* Distant; *Mariekea* Jong & Boer; *Prasia* Stål.

**Comments:** New classifications were proposed to four genera previously classified in the tribe, i.e. *Bafultalna*, *Iruana*, *Murphyalna* and *Sapantanga*. A phylogenetic hypothesis showed Prasiini non-monophyletic including *Iruana* and *Sapantanga*, and both genera were transferred to Hemidictyini (Ruschel & Campos, 2019). However, a new phylogenetic hypothesis was presented, and *Iruana,* *Bafultalna* and *Murphyalna* were transferred to the new tribe, Iruanini (Tettigomyiinae), and a new monogeneric tribe was proposed to accommodate *Sapantanga* (Sapantagini) in Tibicininae (Sanborn et al. 2020).

**Taphurini Distant**

*Dulderana* Distant; *Elachysoma* Torres; *Imbabura* Distant; *Malloryalna* Sanborn; †*Minyscapheus* Poinar, Kritsky & Brown; *Psallodia* Uhler; *Taphura* Stål.

**Comments:** New classifications were proposed to four genera previously classified in the tribe, *Anopercalna*, *Chalumalna*, *Dorachosa*, and *Prosotettix*. *Anopercalna* was transferred to Malagasiini Moulds & Marshall, 2018 by Sanborn (2021) and the subtribe Anopercalnina Boulard, 2008 was then incorrectly synonymized with this tribe. Anopercalnini (Tettigomyiinae) was established based on the Principle of Priority (ICNZ 1999), including *Anopercalna* (Dmitriev & Sanborn, 2023). *Chalumalna* was transferred from Taphurini to Lamotialnini (Sanborn 2025c), *Dorachosa* to Carinetini (Kratzer, 2024), and *Prosotettix* to Selymbriini (Sanborn 2021b). The fossil genus *Minyscapheus* was placed in Taphurini (Moulds, 2018).

**Cicadinae Batsch**

**Antankariini Sanborn, 2021**

*Antankaria* Distant; *Orientafroinsularis* Sanborn.

**Comments:** Antankariini was proposed to include species from Madagascar and the Republic of Seychelles (Africa) (Sanborn, 2021a). Based on the phylogenetic results presented by Marshall et al. (2018) and morphological evidence, Sanborn (2021a) proposed the new tribe and the new genus, *Orientafroinsularis*, to include species of *Chremistica* (Tacuini = Cryptotympanini) from Madagascar, as well transferred *Antankaria* from Tacuini (= Cryptotympanini) to the tribe.

**Arenopsaltriini Moulds, 2018**

*Arenopsaltria* Ashton; *Henicopsaltria* Stål; †*Tithopsaltria* Moulds, Frese & McCurry

Comments: A new fossil genus, *Tithopsaltria*, was proposed from Australia (Moulds, Frese & McCurry, 2022).

**Ayuthiini Moulds, Lee & Marshall**

*Ayuthia* Distant; *Distantalna* Boulard.

**Comments:** The tribe was established to *Ayuthia* and *Distantalna*, previously classified in Tosenini, based in phylogenetic and morphological evidence (Hill et al. 2021).

**Burbungini Moulds**

*Burbunga* Distant; †*Burbungoides* Moulds, Frese & McCurry.

**Comments:** A new fossil genus, *Burbungoides*, was proposed from Australia (Moulds, Frese & McCurry, 2022).

**Cicadini Batsch**

*Cicada* Linnaeus.

**Comments:** The tribe was included in a molecular phylogeny along with 12 related tribes from the Asian continent (Hill et al. 2021).

**Cicadmalleini Boulard & Puissant**

*Cicadmalleus* Boulard & Puissant.

**Comments:** The tribe was sampled in the same phylogeny mentioned in the comment above and recovered in an uncertain relationship with four tribes (Hill et al. 2021).

**Cosmopsaltriini Kato**

*Aceropyga* Duffels; *Brachylobopyga* Duffels; *Cosmopsaltria* Stål; *Diceropyga* Stål; *Dilobopyga* Duffels; *Inflatopyga* Duffels; *Moana* Myers; *Rhadinopyga* Duffels.

**Comments:** The tribe was included in a molecular phylogeny along with 12 related tribes from the Asian continent. Cosmopsaltriini was recovered monophyletic in the genetic tree (Hill et al. 2021).

**Cyclochilini Distant**

*Cyclochila* Amyot & Audinet-Serville.

**Distantadini Orian**

*Distantada* Orian.

**Dundubiini Distant**

*Aola* Distant; *Ayesha* Distant; *Biura* Lee & Sanborn; *Cantata* Lee & Pham; *Champaka* Distant; *Changa* Lee; *Cochleopsaltria* Pham & Constant; *Crassopsaltria* Boulard; *Dundubia* Amyot & Audinet-Serville; *Haphsa* Distant; *Karenia* Distant; Kaphsa Lee; *Khimbya* Distant; *Lethama* Distant; *Macrosemia* Kato; *Megapomponia* Boulard; *Meimuna* Distant; *Minilomia* Lee; *Orientopsaltria* Kato; *Platylomia* Stål; *Sinapsaltria* Kato; *Sinosemia* Matsumura; *Sinotympana* Lee; *Songga* Lee; † *Tymocicada* Becker-Migdisova; *Unipomponia* Lee; *Zaphsa* Lee & Emery.

**Comments:** The fossil genus *Tymocicada* was transferred from Dundubiini to Tacuini (= Cryptotympanini) (Moulds, 2018). *Karenia* was transferred from the monogeneric Sinosenini Boulard based on phylogenetic evidence and Sinosenini was considered a junior synonym of Dundubiini (Hill et al. 2021).

**Durangonini Moulds & Marshall**

*Durangona* Distant.

**Fidicinini Distant**

*Acanthoventris* Ruschel; *Ariasa* Distant; *Beameria* Davis; *Bergalna* Boulard & Martinelli; *Cracenpsaltria* Sanborn; *Diceroprocta* Stål; *Dorisiana* Metcalf; *Elassoneura* Torres; *Fidicina* Amyot & Audinet-Serville; *Fidicinoides* Boulard & Martinelli; *Guyalna* Boulard & Martinelli; *Hemisciera* Amyot & Audinet-Serville; *Hyantia* Stål; *Majeorona* Distant; *Nosoarna* Ruschel & Sanborn; *Nosola* Stål; *Ollanta* Distant; *Orialella* Metcalf; *Pacarina* Distant; *Pompanonia* Boulard; *Prasinosoma* Torres; *Proarna* Stål; *Quesada* Distant; *Rhaeboepelis* Ruschel & Sanborn; *Tympanoterpes* Stål.

**Comments:** A total evidence analysis including some genera of the tribe was published and the new genus, *Acanthoventris*, was proposed (Ruschel et al. 2023). Additionally, other publications proposed new genera to the tribe: *Muraoides* and *Nosoarna* (Sanborn 2018; Ruschel and Sanborn 2021). A new phylogeny of the tribe including molecular data is currently in progress (Ruschel et al. in prep.). *Mura* and *Muraoides* were transferred from Fidicinini to Zammarini based on morphological evidence (Sanborn 2025d).

**Gaeanini Distant**

*Ambragaeana* Chou & Yao; *Balinta* Distant; *Callogaeana* Chou & Yao; *Gaeana* Amyot & Audinet-Serville; *Paratalainga* He; *Sulphogaeana* Chou & Yao; *Talainga* Distant; *Taona* Distant; *Trengganua* Moulton; *Vittagaeana* Moulds, Sarkar, Lee & Marshall.

**Comments:** A molecular phylogeny including Gaeanini was published focusing in 13 tribes with occurrence in the Asian continent (Hill et al. 2021). Gaeanini was recovered non-monophyletic, with *Becquartina* recovered in a different clade and a species of *Tosena* (Tosenini) was recovered closely allied to the sampled genera of Gaeanini in the tree. However, the authors opted only to propose the new genus *Vittagaeana* to the tribe to include two species previously classified in *Tosena* (Tosenini), pointed out the necessity of further studies to the tribe (Hill et al. 2021). Later, a phylogeny of the Gaeanini based on molecular data was published, including genomic data of cicadas obligate symbionts (Wang et al. 2025b). Based on the phylogenetic analysis and morphological evidence, *Becquartina* Kato was transferred from Gaeanini to Leptopsaltriini (Wang et al. 2025).

**Jassopsaltriini Moulds**

*Jassopsaltria* Ashton.

**Kimberpsaltriini Moulds, Marshall, & Popple**

*Kimberpsaltria* Moulds, Marshall & Popple.

**Comments:** The tribe was established to the new genus *Kimberpsaltria* (Moulds et al. 2021).

**Lahugadini Distant**

*Lahugada* Distant.

**Comments:** The tribe was included in a molecular phylogeny and recovered, although with low support, as the most related tribe of Cicadini (Hill et al. 2021).

**Leptopsaltriini Moulton**

*Aetanna* Lee; *Angusta* Wang & Wei; *Becquartina* Kato; *Brevitanna* Lee; *Cabecita* Lee; *Calcagninus* Distant; *Euterpnosia* Matsumura; *Formocicada* Lee & Hayashi; *Formosemia* Matsumura; *Galgoria* Lee; *Indopurana* Lee; *Inthaxara* Distant; *Kalabita* Moulton; *Leptopsaltria* Stål; *Leptosemia* Matsumura; *Manna* Lee & Emery; *Masamia* Lee & Emery; *Maua* Distant; *Metapurana* Lee; *Minipomponia* Boulard; *Miniterpnosia* Lee; *Mosaica* Lee & Emery; *Nabalua* Moulton; *Neocicada* Kato; *Neopurana* Lee & Marshall; *Neoterpnosia* Lee & Emery; *Paranosia* Lee; *Paratanna* Lee; *Philipurana* Lee; *Purana* Distant; *Purapurana* Lee; *Puranoides* Moulton; *Qurana* Lee; *Rustia* Stål; *Sahyaterpnosia* Sadasivan; *Taiwanosemia* Matsumura; *Tanna* Distant; † *Tanyocicada* Moulds; *Versicolora* Wei, Wang, Hayashi, He & Pham; *Vietanna* Lee & Pham; *Yezoterpnosia* Matsumura.

**Comments:** *Gudaba* Distant was shown to be a junior synonym of *Rustia* (Marathe et al. 2018). The fossil genus *Tanyocicada* was proposed for the tribe (Moulds 2020). *Versicolora* was proposed to accommodate two new species, one of them with colour-changing behaviour reported for the first time to Cicadoidea (Wei et al. 2020). In the next year, Leptopsaltriini was included in a molecular phylogeny together with 13 tribes with occurrence in the Asian continent and was recovered polyphyletic (Hill et al. 2021) (see comments in Gaeanini). Based on the results of this study, *Kalabita* Moulton was transferred from Platypleurini to Leptopsaltriini (Hill et al. 2021). *Brevitanna* and *Sahyaterpnosia* was proposed to the tribe based on morphology (Lee 2022; Sadasivan and Sarkar 2023). *Neopurana* was sampled in the phylogeny of Hill et al. (2021) but later placed tentatively in the subtribe Euterpnosiina (Lee et al. 2023). *Vietanna* was proposed based on morphological evidence, and later *Duffelsa* Wang, Jiang & Wei, 2023 was considered a junior synonym of the genus (Pham and Lee 2021; Lee 2023). *Angusta* was described and classified in the tribe based on phylogeny and morphological evidence (Wang and Wei 2024). In a review of *Purana*, four new genera were proposed, *Purapurana*, *Philipurana*, *Metapurana*, and *Indopurana* (Lee 2024a). A posterior molecular phylogeny, focusing on Gaeanini, recovered two species of *Becquartina* Kato as sister-group to the representative clade of Leptopsaltriini, and the genus was transferred from Gaeanini to this tribe (Wang et al. 2025).

**Macrotristriini Moulds**

*Illyria* Moulds; *Macrotristria* Stål; *Mouldspsaltria* Sanborn.

**Comments:** *Mouldspsaltria* was described for the tribe (Sanborn, 2021).

**Oncotympanini Ishihara**

*Mata* Distant; *Neoncotympana* Lee; *Oncotympana* Stål.

**Comments:** The tribe was included in a molecular phylogeny including more 12 tribes with occurrence in the Asian continent. Oncotympanini was recovered sister to *Psithyristria*, part of the sampled Psithyristriini (Hill et al. 2021).

**Platypleurini Schmidt**

*Afzeliada* Boulard; *Albanycada* Villet; *Asianopleura* Zhou, Wang & Wei; *Attenuella* Boulard; *Azanicada* Villet; *Brevisiana* Boulard; *Canualna* Boulard; *Capcicada* Villet; *Dyticopycna* Sanborn; *Eopycna* Sanborn; *Esada* Boulard; *Hainanosemia* Kato; *Hamza* Distant; *Ioba* Distant; *Kalabita* Moulton; *Karscheliana* Boulard; *Koma* Distant; *Kongota* Distant; *Muansa* Distant; *Munza* Distant; *Neoplatypleura* Kato; *Orapa* Distant; *Oxypleura* Amyot & Audinet-Serville; *Planopleura* Lee; *Platypleura* Amyot & Audinet-Serville; *Pycna* Amyot & Audinet-Serville; *Sadaka* Distant; *Sechellalna* Boulard; *Severiana* Boulard; *Soudaniella* Boulard; *Strumosella* Boulard; *Strumoseura* Villet; *Suisha* Kato; *Tigripleura* Lee; *Ugada* Distant; *Umjaba* Distant; *Yanga* Distant.

**Comments:** After Lee (2014) proposed Platypleurini as a junior synonym of Hamzini, an opinion to conserve the usage of the family-group name Platypleurini and its derivative was applied (Marshall et al. 2018a) resulting in Opinion 2466 (2020) followed by Official correction 135 (2021) together conserving Platypleurini. A molecular phylogeny was published for Platypleurini. Based on the results, *Azanicada* Villet was considered a junior synonym of *Platypleura*, and the monogeneric tribe Orapini Boulard a junior synonym of Platypleurini. Consequently, *Orapa* was transferred to Platypleurini (Price et al. 2019). *Pycnoides* and *Eopycna* were proposed to the tribe based on the results of the previously mentioned phylogeny of Platypleurini (Sanborn 2020a). However, later, *Pycnoides* had the name replaced to *Dyticopycna* (Sanborn 2020b). Later, the tribe was included in another molecular phylogeny including 12 tribes related to Cicadini with occurrence in the Asian continent (Hill et al. 2021). *Tugelana* Distant was found to be a junior synonym of *Platypleura* (Villet and Edwards 2021). *Neoplatypleura* had its status revalidated from synonymy with *Platypleura*, and *Planopleura* and *Tigripleura* were proposed to the tribe (Lee 2024b).

**Polyneurini Amyot & Audinet-Serville**

*Angamiana* Distant; *Formotosena* Kato; *Graptopsaltria* Stål; *Parapolyneura* Wang, Hayashi & Wei; *Polyneura* Westwood; *Proretinata* Chou & Yao.

**Comments:** The tribe was included in a molecular phylogeny including 13 tribes with occurrence in the Asian continent. Polyneurini was recovered in a well-supported clade sister to Sonatini. Nonetheless, the phylogenetic relationships recovered among the sampled species of the tribe did not correspond to the currently proposed subdivision into two subtribes (Hill et al. 2021). A more recent phylogeny based on morphological characters and molecular data was published for the tribe, resulting in similar results. The subtribes composition was redefined, and the new genus *Parapolyneura* were proposed. Additionally, *Proretinata* had its status revalidated from synonymy with *Angamiana* (Wang et al. 2024).

**Psaltodini Moulds**

*Anapsaltoda* Ashton; *Neopsaltoda* Distant; *Psaltoda* Stål.

**Psithyristriini Distant**

*Basa* Distant; *Kamalata* Distant; *Onomacritus* Distant; *Ottugia* Lee; *Pomponia* Stål; *Psithyristria* Stål; *Semia* Matsumura; *Terpnosia* Distant; *Trombonia* Lee.

**Comments:** Psithyristriini was recovered polyphyletic in a molecular phylogeny including more 12 tribes with occurrence in the Asian continent (Hill et al. 2021) (see comments in Gaeanini, Leptopsaltriini and Oncotympanini). Sanborn (2025a) proposed the resurrection of Pomponiini, including *Pomponia* Stål, from its synonymy with Psithyristriini based on phylogenetic evidence of Hill et al. (2021) and Wang et al. (2025a). However, Lee (2025) reinstated Pomponiini as a synonym of Psithyristriini, arguing that the phylogenetic analyses in question included members of the *Pomponia linearis* species group instead of the type species of *Pomponia*. According to Lee (2025), the *Pomponia linearis* species group presents morphological differences relative to the type species of *Pomponia*, and form a distinct genus within Psithyristriini.

**Sonatini Lee**

*Hyalessa* China.

**Comments:** There is some confusion over the classification of *Hyalessa*. The genus is currently a senior synonym of *Sonata* Lee classified in Sonatini (Marshall et al. 2018b). Hayashi (2011) synonymized *Sonata* Lee, 2010, type genus of Sonatini, with *Hyalessa* China, 1925, previously classified in Cicadini. The family-group name retains priority (Art. 40.1; ICNZ 1999) and Sonatini remains the name of the tribe even including only *Hyalessa*. A recent phylogeny of Polyneurini included *Hyalessa maculaticollis* (Motschulsky) as outgroup (Wang et al. 2024). Although correctly listed in the material and methods section as a member of Sonatini, the species was erroneously shown in Table 2 of the article as a member of Cicadini (see Wang et al. 2024). The species representing Sonatini was recovered as sister group to Polyneurini, consistent with Hill et al. (2021) results.

**Tacuini Distant**

*Auritibicen* Lee; *Cacama* Distant; †*Camuracicada* Moulds; *Chremistica* Stål; *Cornuplura* Davis; *Cryptotympana* Stål; *Hadoa* Moulds; *Heteropsaltria* Jacobi; *Lyristes* Horváth; *Megatibicen* Sanborn & Heath; *Neotibicen* Hill & Moulds; *Nggeliana* Boulard; *Raiateana* Boulard; *Salvazana* Distant; *Tacua* Amyot & Audinet-Serville.

**Comments:** Tacuini Distant, 1904 was incorrectly established as a junior synonym of Cryptotympanini Handlirsch, 1925 based on phylogenetic analysis and morphological evidence (Marshall et al. 2018b). However, following the Principle of Priority (Art. 23; ICZN, 1999), Tacuini has priority under the Cryptotympanini and was established as the correct taxon name to be used (Dmitriev and Sanborn 2023). The fossil genus *Camuracicada* was proposed to Cryptotympanini (Moulds, 2018). *Antankaria* Distant was removed from the tribe to be part of the new tribe Antankariini proposed by Sanborn (2021a). *Hea* was transferred to the Cicadettini (Cicadettinae) based on morphological evidence (Sanborn, 2023).

**Talcopsaltriini Moulds**

*Talcopsaltria* Moulds.

**Tamasini Moulds**

*Parnkalla* Distant; *Parnquila* Moulds; *Tamasa* Distant.

**Thophini Distant**

*Arunta* Distant; *Thopha* Amyot & Audinet-Serville.

**Tosenini Amyot & Audinet-Serville**

*Tosena* Amyot & Audinet-Serville.

**Comments:** The tribe was recovered polyphyletic in a molecular phylogeny including more 12 tribes with occurrence in the Asian continent (Hill et al. 2021) (see comments in Gaeanini and Leptopsaltriini). Consequently, *Ayuthia* and *Distantalna* were transferred to the new tribe Ayuthiini.

**Zammarini Distant**

*Adusella* Haupt; *Borencona* Davis; *Chinaria* Davis; *Daza* Distant; *Dazollina* Sanborn; *Dyticodopoea* Sanborn; *Heatharia* Sanborn; *Juanaria* Distant; *Miranha* Distant; *Mura* Distant; *Muraoides* Sanborn; *Odopoea* Stål; *Onoralna* Boulard; *Orellana* Distant; *Pachypsaltria* Stål; *Plautilla* Stål; *Pygmaeodopoea* Sanborn; *Procollina* Metcalf; *Uhleroides* Distant; *Zammara* Amyot & Audinet-Serville; *Zammaralna* Boulard & Sueur.

**Comment:** Sanborn (2020c) proposed Plautillini as a junior synonym of Zammarini, and the subtribe Plautillina to include *Plautilla* and *Onoralna*. This change in status of Plautillini was proposed before in a Doctoral thesis, but without recognition by the ICNZ. Kratzer (2024) proposed transferring *Pachypsaltria* to Carinetini (Cicadettinae). However, Sanborn (2025) provided morphological evidence to classify the genus in Zammarini. Four new genera were proposed for the tribe in different publications: *Dazollina, Dyticodopoea, Heatharia,* and *Pygmaeodopoea* (Sanborn 2018, 2020d, 2024b). *Mura* and *Muraoides* were transferred from Fidicinini to Zammarini based on morphological evidence (Sanborn 2025d).

**Derotettiginae Moulds, 2019**

**Derotettigini Moulds**

*Derotettix* Berg.

**Comments:** The new monogeneric subfamily was established based on phylogenetic evidence recovering *Derotettix* as a relict lineage, sister-group of the other subfamilies (Simon et al. 2019).

**Tettigomyiinae Distant**

**Anopercalnini Boulard**

*Anopercalna* Boulard; *Cephalalna* Boulard; *Deremeces* Sanborn; *Ligymolpa* Karsch; *Malagasia* Distant; *Malgachialna* Boulard; *Nyara* Villet; *Quintilia* Stål.

**Comments:** Malagasiini Moulds & Marshall, 2018 was established based on phylogenetic and morphological evidence (Marshall et al. 2018). Later, *Anopercalna* was transferred from Taphurini to the tribe, and Anopercalnina Boulard, 2008 was defined as junior synonym of Malagasiini (Sanborn 2021a). Additionally, *Cephalalna* was transferred to the tribe and *Deremeces* was described for the tribe (Sanborn, 2021a). However, the name Anopercalnina has priority under Malagasiini (Art. 23; ICZN, 1999), and the error was corrected making Malagasiini junior synonym of Anopercalnini (Dmitriev & Sanborn, 2023).

**Hovanini** **Sanborn, Marshall & Moulds**

*Hovana* Distant.

**Comments:** *Hovana*, before classified in Hemidictyini (Marshall et al., 2018; Ruschel and Campos, 2019), is assign to the monogeneric tribe *Hovanini* (Tettigomyiinae) based on molecular phyologenetic analysis and morphological evidence (Sanborn et al. 2020).

**Iruanini Boulard**

*Bafutalna* Boulard; *Murphyalna* Boulard; *Iruana* Distant; *Lacetas* Karsch.

**Comments:** See Prasiini section. Based on the results of the phylogenetic analysis, *Lacetas* was transferred from Hemidictyini to Iruanini, making Lacetasini Moulds & Marshall, 2018 a junior synonym of Iruanini Boulard, 1993 (Art. 23; ICZN, 1999) (Sanborn et al., 2020).

**Tettigomyiini Distant**

*Bavea* Distant; *Gazuma* Distant; *Paectira* Karsch; *Spoerryana* Boulard; *Stagea* Villet; *Stagira* Stål; *Tettigomyia* Amyot & Audinet-Serville; *Xosopsaltria* Kirkaldy.

**Ydiellini Boulard**

*Maroboduus* Distant; *Nablistes* Karsch.

**Tibicininae Distant**

**Aragualnini Sanborn**

*Aragualna* Champanhet, Boulard & Gaiani.

**Comments:** The tribe, before classified in Cicadettinae, was transferred to Tibicininae based on molecular phylogeny and morphological evidence (Owen et al. 2022).

**Chilecicadini Sanborn**

*Chilecicada* Sanborn.

**Hemidictyini Distant**

*Hemidictya* Burmeister.

**Comments:** Hemidictyini, previously classified in Cicadettinae, was tentatively transferred to Tettigomyiinae based on phylogenetic evidence (Ruschel and Campos, 2019). Later, a new hypothesis was proposed, and the tribe is currently classified in Tibicininae (Sanborn et al. 2020).

**Platypediini Kato**

*Neoplatypedia* Davis; *Platypedia* Uhler.

**Selymbriini Moulds & Marshall**

*Prosotettix* Jacobi; *Selymbria* Stål; *Striduloselymbria* Sanborn.

**Comments:** *Prosotettix* was transferred from Taphurini (Cicadettinae) to the tribe (Sanborn 2021b), and *Striduloselymbria* was described for the tribe (Sanborn 2024c).

**Tettigadini Distant**

*Acuticephala* Torres; *Alarcta* Torres; *Babras* Jacobi; *Calliopsida* Torres; *Chonosia* Distant; *Coata* Distant; *Mendozana* Distant; *Psephenotettix* Torres; *Tettigades* Amyot & Audinet-Serville; *Tettigotoma* Torres; *Torrescada* Sanborn & Heath.

**Tibicinini Distant**

*Chlorocanta* Chatfield-Taylor; *Clidophleps* Van Duzee; †*Davispia* Cooper; *Gibbocicada* Ruschel; *Hewlettia* Smeds; †*Lithocicada* Cockerell; *Okanagana* Distant; *Okanagodes* Davis; *Paharia* Distant; *Subpsaltria* Chen; *Subtibicina* Lee; *Tibicina* Kolenati; *Tibicinoides* Distant.

**Comments:** Three genera are described, and two fossil genera are placed in the tribe (Moulds 2018; Ruschel 2018; Cole et al. 2023).

**Fossil genera with only family or subfamily classification:** *Burmacicada* Poinar & Kritsky; *Dominicicada* Poinar & Kritsky; *Feoichnus* Krause, Bown, Bellosi & Genise; *Fonsecacicada* Martins-Neto & Mendes; *Kintusamo* Szwedo.
